# TESSA: A unified model to detect trajectory-preserved and spatially-variable genes in spatial transcriptomics

**DOI:** 10.1101/2025.09.06.674654

**Authors:** Yuesong Wu, Haohao Su, Nina Steele, Yuying Xie, Yuehua Cui

## Abstract

Identifying spatially variable genes (SVGs) has been an essential task in spatial transcriptomics. In addition to SVGs detection, there are genes exhibiting expression patterns that are associated with cellular developmental stages or lineage fates across a tissue section. Identifying such genes could provide novel insights into tumor metastasis. Here, we introduce a unified statistical model, termed TrajEctory-preServed and SpAtially-variable gene detection (TESSA), to detect both types of genes. Moreover, we propose a novel strategy to address the inherent double-dipping issue commonly encountered when assessing temporal gene effects in transcriptomics analysis. We demonstrate the testing performance through extensive simulation studies and real applications to several publicly available datasets. Downstream analyses further highlight the potential of our method in identifying genes associated with tumor progression and enhancing spatial domain detection.

## 1 Introduction

Spatial transcriptomics (ST) has revolutionized our understanding of gene expression within the intact tissue context. By preserving spatial information, it enables researchers to uncover tissue anatomical architecture, local microenvironments, and region-specific gene regulation. Spot-based ST platforms measure expression levels for the entire transcriptomic gene panel (e.g., 10x Visium), while cellular-resolution technologies profile up to thousands of genes per cell (e.g., 10x Xenium)[1]. However, due to the limited gene panel that can be measured by single-cell resolution technology, spot-level ST data are still predominantly used in which detecting spatially variable genes (SVGs) remains a major task[2]. SVGs exhibit distinct expression patterns across tissue regions, implicating their involvement in region-specific biological processes[3]. SVG detection has been proven crucial for constructing low-dimensional spot/cell embeddings[4], detecting spatial domains[5], and spatial boundaries[6]. It also enables the identification of domain-specific markers[7] and cell type-specific spatially variable markers[8]. However, most SVG detection methods focus exclusively on spatial variability, without explicitly modeling the dynamic transcriptional changes that occur along developmental stages or lineage fates. For example, biological processes such as embryonic development, cell differentiation, and tumor progression are not only spatially heterogeneous but also temporally dynamic. Thus, there is a pressing need to develop a unified statistical method that can infer both temporally and spatially variable genes in ST analysis, which we refer to as TSVGs.

The temporal information is well-defined in the scRNA-seq field as the concept of pseudotime[9], which reconstructs developmental trajectories from snapshot data even though the actual physical time of cell state transitions is unobservable. Based on gene expression profiles of individual cells, algorithms such as Monocle[10], Slingshot[11], and DPT[12] can reconstruct dynamic trajectories or pseudotime in high-dimensional gene expression space, which provides an inferred ordering of cells within the same tissue compartment. For example, for a specimen of colon tissue containing cells in different cell states, the inferred pseudotime trajectory reveals a continuum from intestinal stem cells to transit-amplifying progenitors and finally to fully differentiated absorptive and secretory epithelial cells[13]. To further explore the biological significance of these trajectories, trajectory-based differential expression analysis methods have been developed to detect temporally variable genes (TVGs). For example, approaches such as TradeSeq[14], TSCAN[15], and PseudotimeDE[16] were developed to identify genes whose expression patterns are associated with lineage transitions.

When a meaningful pseudotime is extended to ST context, even if the pseudotime is inferred purely from gene expression data, it often aligns with a certain non-random spatial structure (Fig. 1; Fig. S6b; Fig. S2b). As a result, TVGs whose expression levels change along pseudotime, also tend to display certain non-random spatial patterns (Fig. 1c2-3). However, standard SVG detection algorithms often fail to identify these TVGs because they rely on direct spatial variability rather than the underlying temporal dynamics. This highlights a key discrepancy between temporal and spatial signals: TVGs may exhibit spatial structure that is not fully captured by conventional SVG detection methods. Currently, there remains a lack of dedicated TVG detection methods tailored for ST context, further complicated by a statistical challenge known as the double dipping issue.

**Fig. 1:**
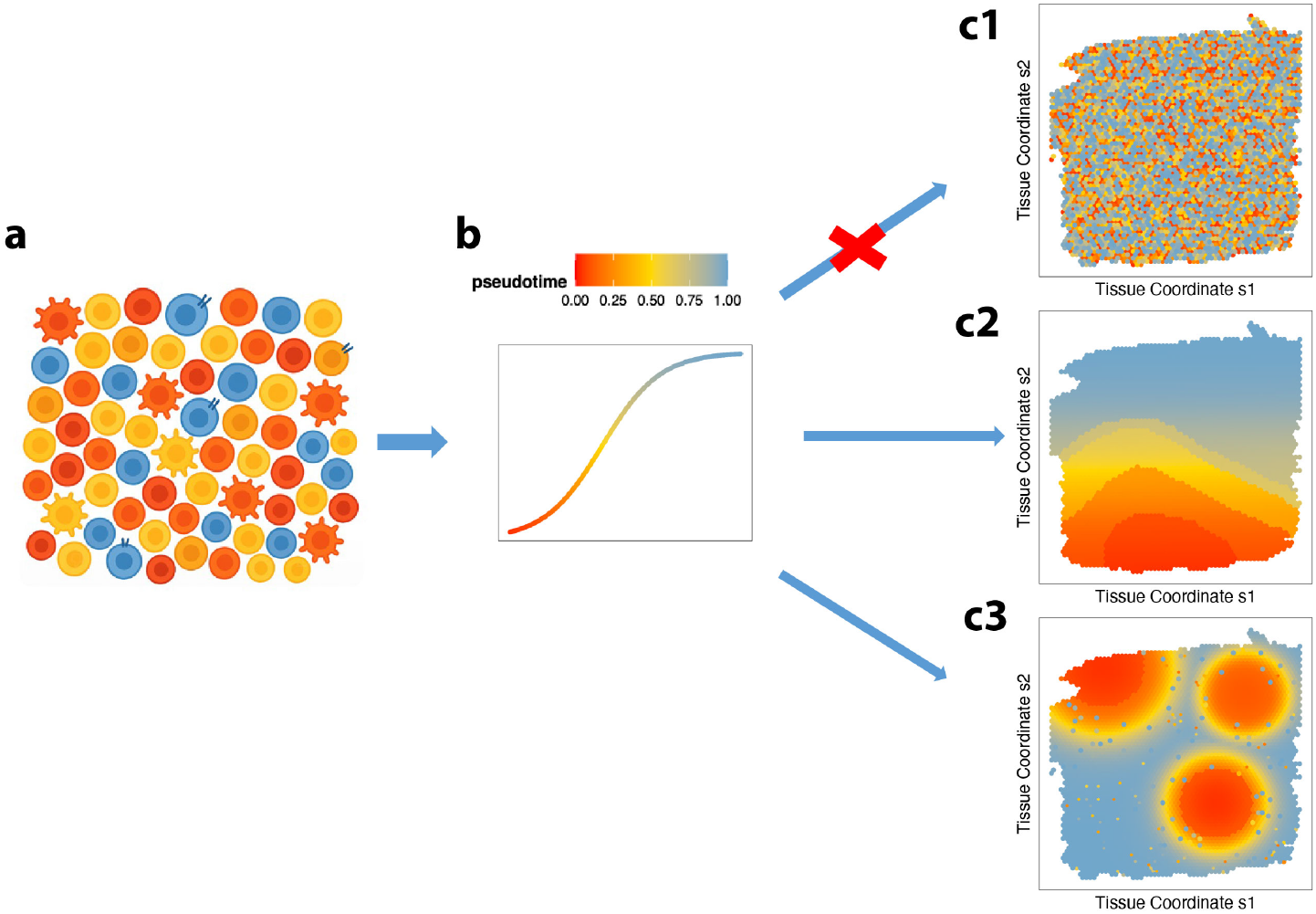
Motivating Toy Example. **a**. Toy example illustrating a bunch of single cells without spatial context. **b**. Cells are sorted by pseudotime order, where pseudotime is inferred solely based on gene expression profiles. **c**. When mapping single cells onto the spatial layout, the cells may exhibit randomly scattered patterns, as seen in c1, or spatially organized patterns, as shown in c2 and c3. If a meaningful pseudotime is present in panel b, the pattern observed in c1 is unlikely to arise.

**Fig. 2:**
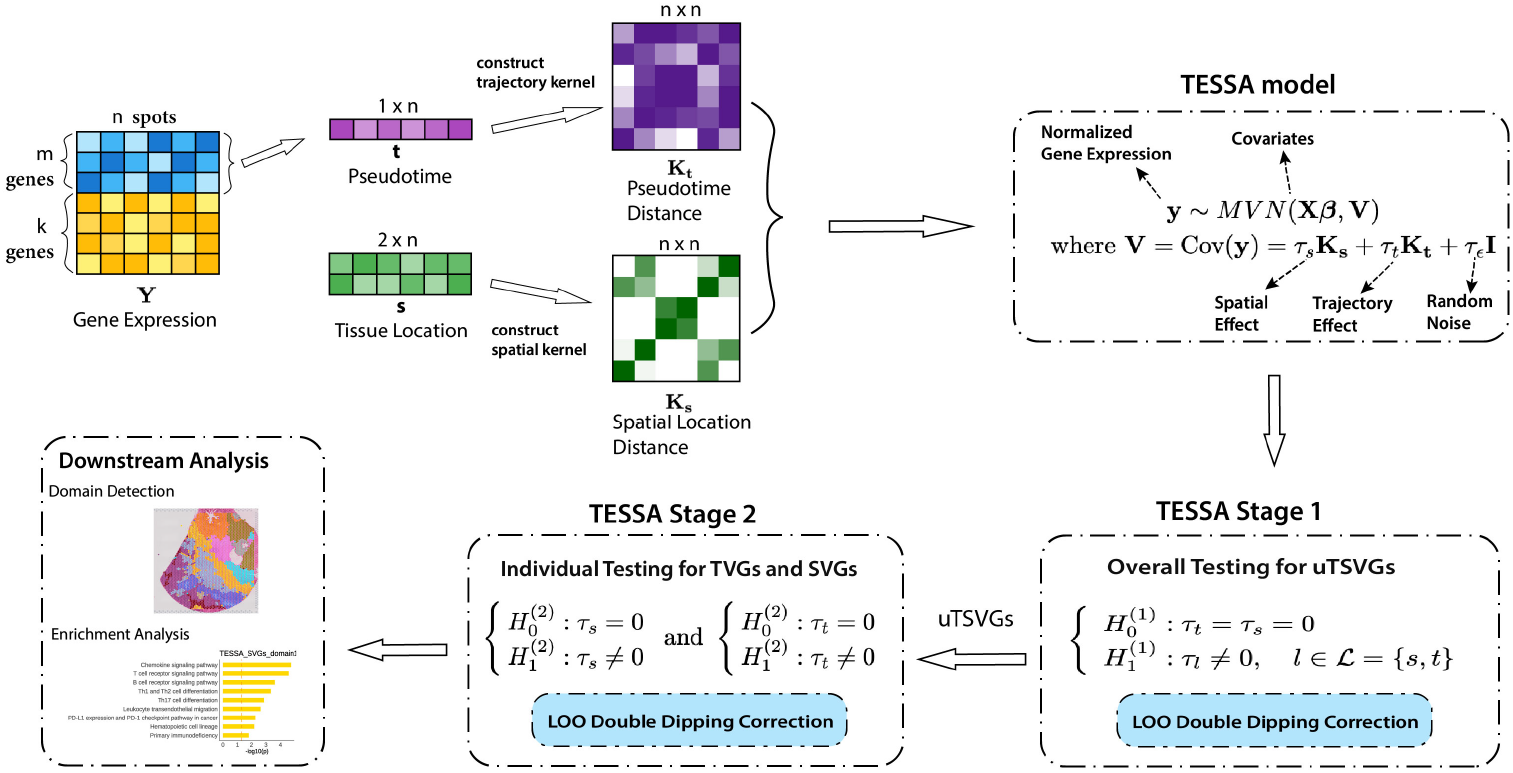
Overview of TESSA. TESSA is a unified statistical model designed to detect both temporally variable genes(TVGs) and spatially variable genes (SVGs) in spatial transcriptomics. Utilizing trajectory information obtained from pseudotime inference, TESSA links gene expression patterns with temporal and spatial kernel matrices through a linear mixed effect model incorporating trajectory and spatial random effects. In stage 1, both TVGs and SVGs are identified through an overall test, which determines if both temporal and spatial variance components are non-zero. Genes passing this unified test are termed as unified temporal and saptial variable genes(uTSVGs). In stage 2, for each variance componet of interest, TVGs or SVGs are determined from the uTSVGs by conducting individual tests to check if the corresponding temporal or spatial variance component is non-zero. Downstream analysis of TESSA includes spatial domain detection using uTSVGs and TVGs gene set enrichment analysis.

Double dipping is a notorious issue in gene testing, arising when latent variables, such as pseudotime or clusters, are used both in the estimation process and in subsequent hypothesis testing [17, 18, 16, 19, 20]. This is because the same data is used twice for both pseudotime estimation and testing, leading to spurious correlations and consequently seriously inflated type I errors. Specifically, in pseudotime analysis, the true, or say the oracle, pseudotime is generally unknown, so the same genes are used twice for both pseudotime estimation and for testing the significant pseudotime effect. Countsplit[17], DataThinning[18], and pseudotimeDE[16] are proposed to address the pseudotime-related double dipping issue for scRNA-seq data. However, their performance on ST analysis has theoretical and practical limitations (see details in the simulation and real data analysis results).

In this work, we propose a simple solution, Leave-One-Out (LOO), to address the double-dipping issue when inferring TVGs in the spatial context in addition to SVG detection. Our method enables the detection of pseudotime-associated genes with effective type I error control while maintaining reasonable scalability. This serves as a necessary statistical foundation for building a unified model for the detection of TrajEctory-preServed and SpAtially-variable genes in spatial transcriptomics (TESSA), with a two-stage hypothesis testing procedure, to identify unified temporally and spatially variable genes (uTSVGs) and to further distinguish them into TVGs and SVGs. By incorporating temporal information, we are able to disentangle key features into temporally and spatially variable genes. These components provide complementary insights into the underlying biology and offer evidence to enhance downstream domain-specific analyses and provide additional biological insight into developmental progression, such as tumor progression, tumor-immune microenvironment, and cortex region development.

## 2 Results

### 2.1 Method overview

The schematic representation of TESSA is displayed in Fig.2. The technical details are provided in the Methods section and Supplementary File Details of TESSA. Specifically, we developed a two-stage testing procedure for TESSA to detect TSVGs: 1) Stage 1 test: the TESSA overall test to identify unified temporally and spatially variable genes (uTSVGs), referring to statistically significant genes that might exhibit at least one of spatial and temporal effects; and 2) Stage 2 test: the two TESSA individual tests to dissect whether a uTSVG is either a TVG or an SVG, respectively. For each stage, we applied our LOO correction to the significant uTSVGs or TVGs to avoid false positives due to double dipping. We demonstrate the benefits of TESSA through downstream analysis, including domain detection and functional enrichment analysis about tumor progression and cortex development.

### 2.2 Simulations

We designed simulation studies to evaluate the performance of TESSA in dissecting temporal and spatial effects, as well as type I error control and detection power with the LOO strategy. We investigated the performance under two data generation scenarios to evaluate the robustness on normalized gene expression data and raw count data. Fig.9 displays the data generation of the Negative Binomial(NB) scenario and Fig.S1a,b,c displays the Normal data generation scenario. Under each scenario, we designed two sets of simulations to assess the performance of the twostage testing framework and compared it with existing methods in SVG detection and double-dipping correction in spatial transcriptomics data. As TESSA relies on pseudotime information to detect uTSVGs and TVGs, we tested TESSA using both the true pseudotime and estimated pseudotime by slingshot[11], denoted as TESSA-oracle and TESSA-slingshot, respectively (see pseudotime estimation details in Supplementary File Pseudotime Estimation). We also evaluated the false positive control of TESSA using the proposed LOO strategy (TESSA-slingshot-LOO) and the Countsplit strategy[17] (TESSA-slingshot-countsplit).

#### 2.2.1 Performance of Stage 1 Overall Test

The first set of simulations evaluated the performance of the TESSA stage 1 overall test for TSVGs detection, the significant genes, termed as unified TSVGs (uTSVGs), refer to genes that has at least one of temporal or spatial effect. We compared it with SPARK gaussian version[21] with 10 kernels for SVG detection and with Countsplit[17] for double dipping correction. We presented the results with the Negative Binomial data generation scenario here. Normal scenario has similar results which are given in Supplementary File 2. We applied VSTNormalization[21] to transform the raw count gene expression data to cotinuous data, to satisfy the model structure of TESSA.

Under the null setting, where all the genes with neither spatial nor temporal patterns, TESSA-oracle effectively controlled the type I error under different data dispersions (Fig.3a; dispersion = 0.7, false positive rate (FP) = 0.0458; dispersion = 1, FP = 0.0514; dispersion = 1.5, FP = 0.0511). However, when directly using the estimated pseudotime, TESSA-slingshot has serious inflated false positives because of double dipping (dispersion = 0.7, FP = 0.2253; dispersion = 1, FP = 0.2201; dispersion = 1.5, FP = 0.2232). While implementing the countsplit strategy, i.e., TESSA-slingshot-countsplit, reduces the inflation, it still cannot control the false positive rate well. Besides, as data dispersion decreases (i.e., variance increases), the efficacy of double dipping correction reduces (dispersion = 0.7, FP = 0.1126; dispersion = 1, FP = 0.0865; dispersion = 1.5, FP = 0.065). On the other hand, TESSA-slingshot-LOO still produced testing results comparable to TESSA-oracle, indicating that LOO is an effective strategy to control type I error due to double dipping (dispersion = 0.7, FP = 0.0509; dispersion = 1, FP = 0.0479; dispersion = 1.5, FP = 0.0522). SPARK only relies on location information, so it does not have the double dipping issue (dispersion = 0.7, FP = 0.0537; dispersion = 1, FP = 0.0509; dispersion = 1.5, FP = 0.0529).

Under the alternative hypothesis setting, we assessed the method’s performance to distinguish true TSVGs from noise genes. Here, the TSVG set comprises genes exhibiting a temporal pattern, a spatial pattern, or both, whereas noise genes (also termed as null genes) were simulated to be independent of any spatial or temporal structure. Performance was quantified by the AUC-ROC (the Area Under the Receiver Operating Characteristic Curve), as a comprehensive metric which jointly evaluates sensitivity (true-positive rate or TPR) and specificity (true-negative rate or TNR). A higher AUC-ROC value indicates that detected uTSVGs closely match the ground-truth TSVGs and that detected noise genes align with the true noise set.

TESSA-oracle, which uses ground-truth pseudotime, achieved the highest AUC-ROC and statistical power (light blue curve in Fig. 4a, b, c). However, in real applications, the oracle pseudotime is generally unknown. As expected, all other methods, relying on estimated pseudotime or spatial information alone, exhibited reduced power. Interestingly, even though the power of TESSA-slingshot (purple curve) demonstrated higher power than TESSA-slingshot-LOO (blue curve) (Fig. 4c), its overall accuracy in distinguishing true TSVGs from noise genes was lower (Fig. 4a). This suggests that TESSA-slingshot misidentified some spurious noise genes that contribute to pseudotime estimation as uTSVGs, and the false positive rate might outweigh the true positive rate in some settings. In contrast, TESSA-slingshot-LOO avoids this issue by excluding the signature gene from pseudotime estimation when testing it with the LOO strategy. As a result, despite having slightly lower power, TESSA-slingshot-LOO achieves better overall classification accuracy by reducing false positives. Additionally, the power of both TESSA-slingshot-countsplit (olive green curve) and SPARK (yellow curve) is low in the detection of TSVGs (Fig. 4c). Power loss is inevitable for Countsplit[17] as its data-splitting strategy inevitably weakens signal strength by splitting the gene expression levels into halves. While SPARK can detect spatially variable signals, it is not designed to capture transcriptional dynamics along developmental trajectories, which limits its statistical power relative to TESSA for detecting true TVGs as part of true TSVGs. For the power to detect TVGs (Fig. 4d), the countsplit strategy also leads to lower power compared to the LOO strategy, while SPARK has the lowest power as it is not designed to capture such a spatial pattern.

#### 2.2.2 Performance of Stage 2 Individual Test

The second set of simulations evaluated the performance of the TESSA stage 2 individual test for TVG and SVG detection separately. We compared it with Countsplit[17] for TVG double-dipping correction.

Under the null setting containing only noise genes, we observed that when testing for SVGs, both TESSA-oracle and TESSA-slingshot effectively controlled the type I error across different levels of data dispersion, as no double dipping occurs in this context (Fig. 3c). However, when testing for TVGs, TESSA-slingshot showed inflated type I error (dispersion = 0.7, FP = 0.1395; dispersion = 1, FP = 0.147; dispersion = 1.5, FP = 0.137), while TESSA-slingshot-LOO achieved good type I error control (dispersion = 0.7, FP = 0.052; dispersion = 1, FP = 0.063; dispersion = 1.5, FP = 0.0495), comparable to TESSA-oracle (dispersion = 0.7, FP = 0.045; dispersion = 1, FP = 0.051; dispersion = 1.5, FP = 0.052) (Fig. 3b). Consistent with the first simulation study, TESSA-slingshot-countsplit partially alleviated false positives but failed to fully control type I error, especially under varying data dispersion levels (dispersion = 0.7, FP = 0.083; dispersion = 1, FP = 0.071; dispersion = 1.5, FP = 0.0665).

**Fig. 3:**
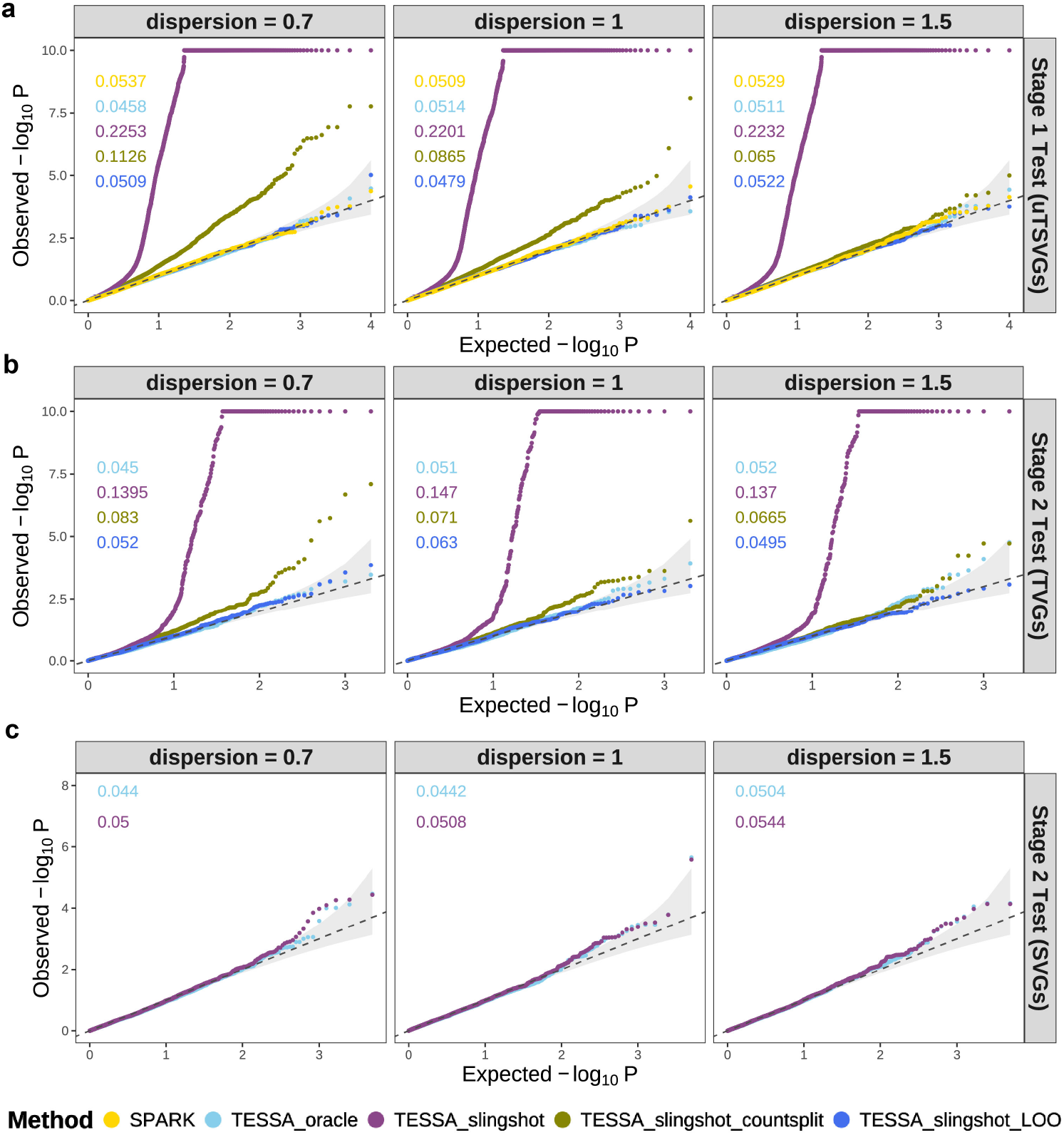
Null Case Results in the Negative Binomial Simulation Scenario. The Q-Q plots of the observed − log_10_ *P* against the expected − log_10_ *P* for different methods are displayed across various dispersion parameters, where P represents the p-value. False positive rates (FP) for each method are displayed in the top-left corner of each panel, with colors corresponding to the legend. The light gray region represents the 95% confidence band, illustrating the range in which 95% of points are expected to fall under the null hypothesis, assuming the p-values follow the expected uniform distribution. **a**. Null case result of simulation 1. Stage 1 Overall Test for TSVG detection; **b**. Null case result of simulation 2. Stage 2 Individual Test for TVG detection; **c**. Null case result of simulation 2. Stage 2 Individual Test for SVG detection.

**Fig. 4:**
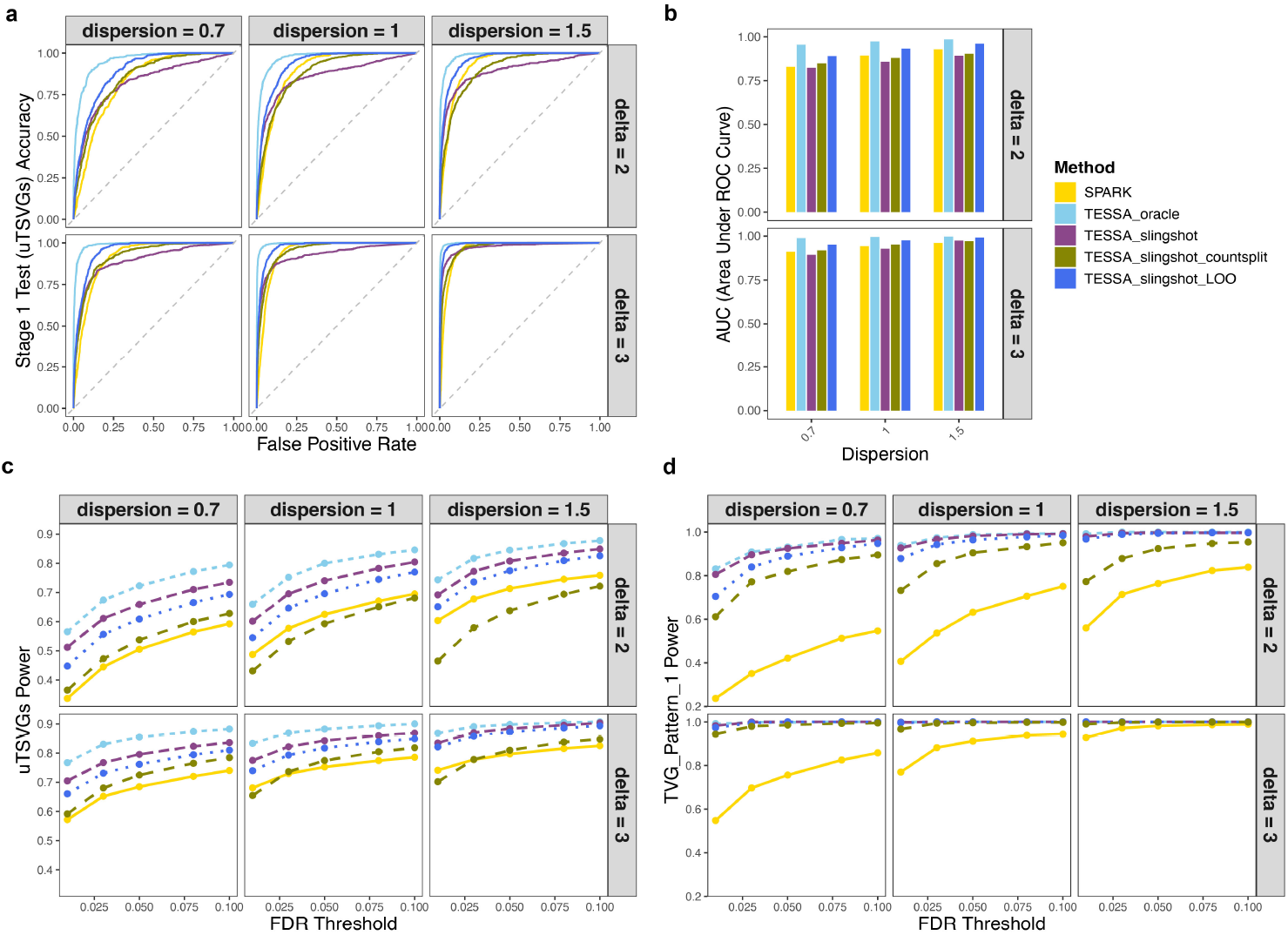
Alternative Case Results of Stage 1 Test in the Negative Binomial Scenario. **a**. The ROC plots of TSVG detection accuracy for different methods displayed across various dispersion parameters and different signal strengths. The true labels are either TSVG or noise (non-TSVG); the x-axis represents the false positive rate and the y-axis represents the true positive rate. **b**. The barplot of Area Under ROC Curve (AUC) for different methods displayed across various dispersion parameters and different signal strengths, corresponding to the ROC curves in plot (a). **c**. The power plots of TSVG detection across a range of FDR levels for different methods, considering various dispersion parameters and signal strengths. **d**. The power plots of TVG detection for data generated with the TVG pattern 1 setting, across a range of FDR levels for different methods, considering various dispersion parameters and signal strengths. SPARK is not designed for TVG detection but still picks up some power.

Under the alternative setting, we compared these methods in terms of their ability to distinguish SVG (or TVG) from null genes, measured by AUC-ROC values. For TVG detection (Fig. 5a,b), TESSA-oracle achieved the highest accuracy, followed by TESSA-slingshot-LOO. As seen in the first simulation set, although TESSA-slingshot achieved higher statistical power, its overall accuracy was low due to increased false positives caused by the double dipping. In contrast, TESSA-slingshot-LOO, by excluding the gene from pseudotime estimation when testing it, achieved better overall classification accuracy. TESSA-slingshot-countsplit showed the lowest power and accuracy, likely due to signal attenuation from the data splitting strategy inherent in Countsplit[17]. Under the two TVG power settings (Fig. 5c,d), TESSA-slingshot-LOO achieved better power than TESSA-slingshot-countsplit. Though TESSA-slingshot had higher power than TESSA-slingshot-LOO, it is likely due to high false positives (3b). For SVG detection (5e,f), there is no need to implement the LOO strategy and TESSA-slingshot achieved nearly the same power as TESSA-oracle, showcasing the robust performance of TESSA for SVG detection.

**Fig. 5:**
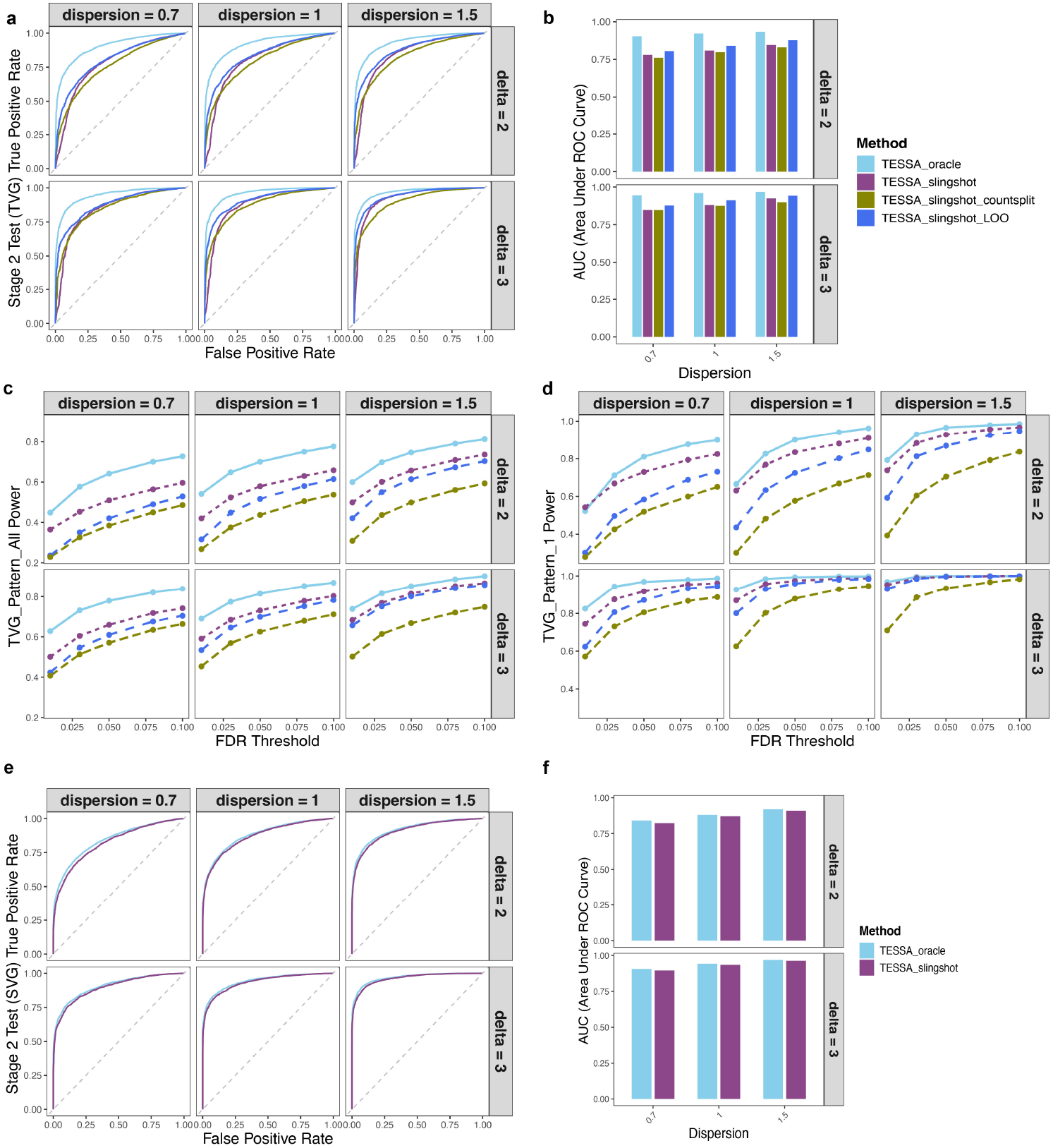
Alternative Case Results of Stage 2 Test in the Negative Binomial Scenario. **a**. The ROC plot of TVG detection accuracy for different methods displayed across various dispersion parameters and different TVG signal strengths (*δ* = 2 and 3). The true labels are either TVGs or non-TVGs, the x-axis represents the false positive rate, and the y-axis represents the true positive rate. **b**. The barplot of Area Under ROC Curve (AUC) corresponding to the ROC curves in **a. c**. The plot of testing power for all TVGs across a range of FDR levels for different methods, considering various dispersion parameters and signal strengths. **d**. The plot of testing power for TVG generated for TVG pattern 1 setting, across a range of FDR levels for different methods, considering various dispersion parameters and signal strengths. **e**. The ROC plot of SVG detection accuracy. The true labels are either SVGs or non-SVGs. **f**. The barplot of Area Under ROC Curve (AUC) corresponding to the ROC curves in plot **e**.

### 2.3 Real Data Analysis

We applied TESSA to three ST datasets and compared its performance with other methods shown in the simulation study.

#### 2.3.1 Human Pancreatic Ductal Adenocarcinoma ST Analysis

We analyzed a 10x Visium spatial transcriptomics dataset from human pancreatic ductal adenocarcinoma (PDAC)[22]. PDAC is a highly aggressive cancer marked by complex tumor-stroma interactions and resistance to therapy. The progression from normal pancreatic regions, such as acinar, to PaNIN (Pancreatic Intraepithelial Neoplasia), to ductal tumor and stroma, including cancer-associated fibroblasts (CAFs), is central to PDAC development. We aim to assess whether integrating pseudotime-related variation through TVGs provides biologically meaningful and complementary information to traditional SVG detection.

We focused our analysis on sample S2A, which contains 3,142 spots and 17,943 genes. After filtering out low-expressed genes, those expressed in fewer than 10% of spots, 11,115 genes remained. Pseudotime was inferred using Slingshot[11] based on 31 curated genes (see Supplementary File Pseudotime Estimation) associated with PDAC progression, resulting in two lineages that initiate from PaNIN regions (Fig. S2b,d). Using the TESSA Stage 1 overall test, we identified 6,956 uTSVGs at an FDR threshold of 0.05 followed by the Benjamini–Yekutieli correction. For comparison, SPARK identified 7,205 SVGs, with 6,309 overlapping with the uTSVGs. TESSA lineage-specific individual tests further identified 3,632 lineage 1 TVGs, 4,019 lineage 2 TVGs, and 5,435 SVGs (Fig. 6d).

**Fig. 6:**
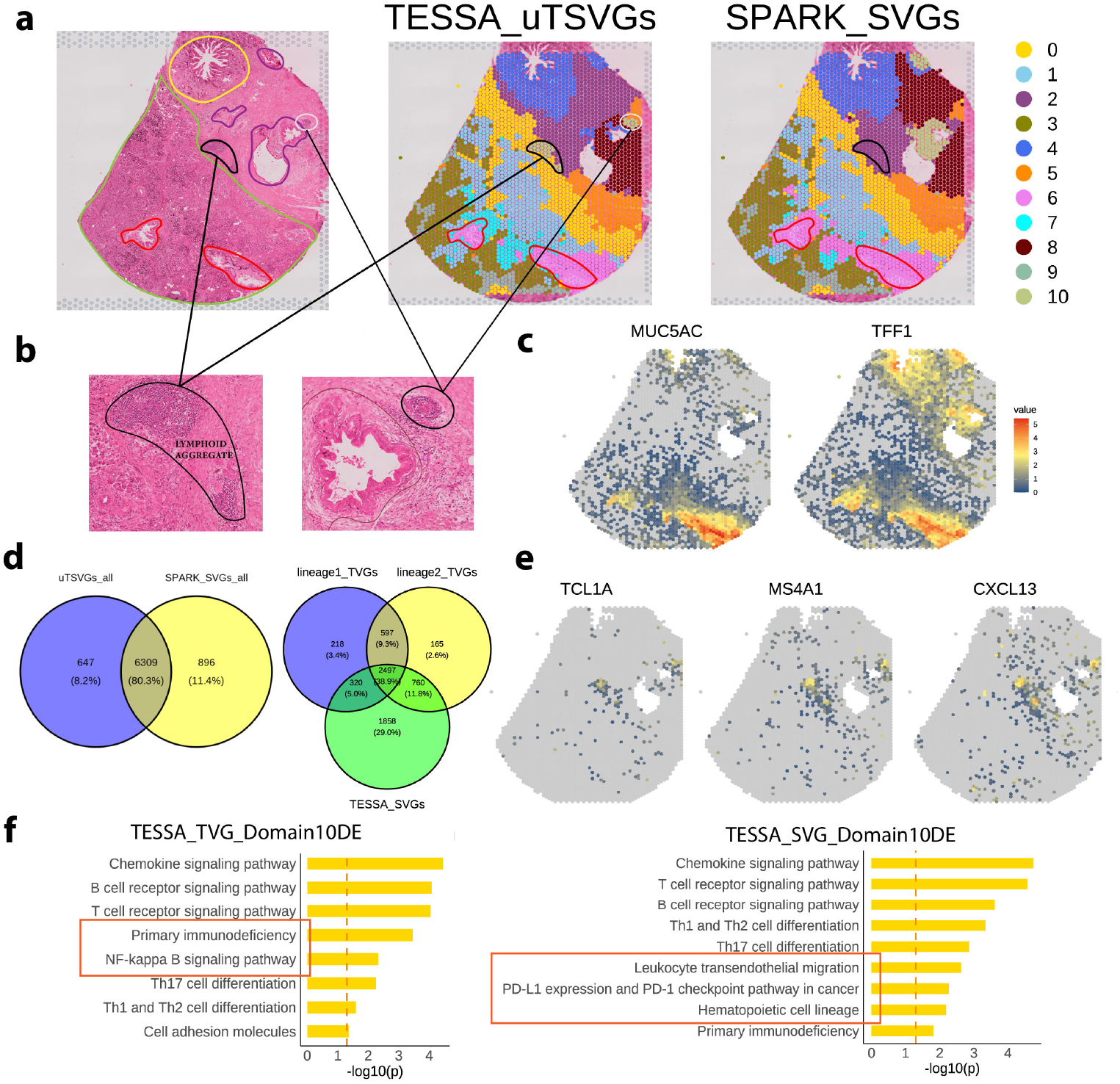
Results of Pancreatic Ductal Adenocarcinoma ST Data Analysis. **a**. Left: Pathologist’s rough annotation as ground truth. Yellow as normal ductal region, purple as tumor ductal region, green as acinar region, red as PaNIN region, and black as lymphoid aggregation region. Middle: SeuratPCA domains based on TESSA TSVGs. Right: SeuratPCA domains based on SPARK SVGs. **b**. Zoomed in H&E image shows that TESSA TSVGs domain 10 is aligned well with the lymphoid aggregation area. **c**. Spatial pattern plots of log-normalized gene expression for PaNIN marker gene *MUC5AC* and pancreatic tumor marker gene *TFF1*. **d**. The left Venn diagram shows the logical relationship between sets of genes identified by TESSA and SPARK. The right diagram shows the logical relationship between sets of lineage-specific TVGs and SVGs identified by TESSA stage 2 tests. **e**. Spatial pattern plots of log-normalized gene expression for TESSA TSVGs domain 10 differentially expressed genes, including B cell marker genes *TCL1A, MS4A1*, and lymphoid structures *CXCL13*. **f**. The KEGG gene set enrichment analysis results for TVGs and SVGs detected by TESSA stage 2 tests, which are also differentially expressed in domain 10. Left: TVGs enriched in immune activation pathways. Right: SVGs enriched in immune regulation pathways.

To evaluate whether incorporating temporal information improves downstream analysis, we compared domain detection performance using uTSVGs from TESSA and SVGs from SPARK. We applied both SeuratPCA[23] and SpatialPCA[4] with default settings to conduct domain detection, and the evaluation referred to the rough pathologist annotations (Fig. 6a left plot). Domain results from spatialPCA based on TESSA uTSVGs and SPARK SVGs are generally similar but fail to capture finegrained morphological structures and tissue heterogeneity (Fig. S3e). Therefore, for clearer comparison and demonstration, we use the domain results from SeuratPCA with 11 clusters (Fig. 6a middle and right plots). Results using different numbers of clusters are provided in Fig. S3a,b.

##### uTSVGs captured PaNIN and lymphoid aggregation domain

Domain results from SeuratPCA based on uTSVGs exhibited finer resolution and better visual alignment with known histological structures. For example, domain 6 more precisely captured the PaNIN region compared to results based on SPARK derived SVGs, which tended to over-merge adjacent areas (Fig. 6a). *MUC5AC* and *TFF1* are both markers of “classical” subtype identity of PDAC cells, but will also mark PanIN lesions, since PanINs largely acquire a classical gene program. The expression of both *MUC5AC* and *TFF1* aligned well with pathologist annotations of the PaNIN region (Fig. 6c), closely matching the shape of domain 6 identified using TESSA uTSVGs.

Notably, domain 10 (Fig. 6b, left), corresponding to lymphoid aggregation, was only detected when using uTSVGs. SPARK failed to isolate this small but biologically meaningful region, instead merging it into domain 2 (Fig. 6a, right). While domain 10 does not exactly match the rough manual annotation, its biological relevance is supported by strong expression of B cell markers (*CR2, MS4A1, CXCL13* ; Fig. 6e). *CXCL13* notably is very important for maturation and infiltration of B cells, and also even the formation of tertiary lymphoid aggregates[24]. The spatial expression pattern of *CXCL13* strongly overlaps with domain 10 (Fig. 6a) and histological features visible in the H&E image (Fig. 6b), validating the improved domain detection resolution achieved with TSVGs by TESSA.

In practical applications, it is common to use only the top-ranked genes for domain analysis rather than the full list of detected uTSVGs or SPARK SVGs [25]. To evaluate this, we compared domain results based on the top 2,000 lineage-specific uTSVGs and an equal number of SPARK-derived SVGs. The results were largely consistent with those obtained using all significant genes, with the top uTSVGs still successfully capturing both domain 6 and domain 10 (see Fig. S3c,d).

##### TVGs captured pseudotemporal regulatory programs beyond spatial signatures

To assess the biological relevance of TVGs, we performed Reactome pathway enrichment analysis using lineage-specific unique TVGs and SVGs (Fig. S2e). Unique TVGs refers to genes that are identified as TVGs but not SVGs, vice versa for unique SVGs. We focus on these non-overlapping TVGs and SVGs, to highlight their distinct functional roles. In lineage 1, TVGs were enriched in pathways such as Rho GTPase signaling, vesicle-mediated transport, and the VEGFA–VEGFR2 signaling axis. These pathways are often associated with cellular movement, intracellular trafficking, and vascular remodeling, suggesting that TVGs in this lineage may reflect dynamic changes in the tumor microenvironment (TME) over pseudotime. In contrast, SVGs from the same lineage-specific analysis were enriched for receptor tyrosine kinase (RTK) signaling, EGFR signaling, and interleukin-mediated pathways, indicating spatially localized immune-tumor interactions and signaling gradients. In lineage 2, which includes more acinar-like and less CAF-rich regions, TVGs were enriched in pathways related to programmed cell death and cell cycle regulation. This suggests a pseudotemporal trend of decreasing proliferation or increased apoptosis. Meanwhile, SVGs were enriched for immune surveillance and regulatory pathways such as MHC class I antigen presentation, Toll-like receptor signaling, and SUMOylation, reflecting the spatial organization of immune activity in these tissues. These results support the idea that TVGs and SVGs capture complementary biological information. While SVGs tend to highlight spatially distinct functional zones, TVGs may better capture gradual regulatory shifts or transitions over pseudotime. Notably, the enrichment of TVGs in coherent and biologically relevant pathways indicates that these TVGs are not just statistical artifacts, but represent meaningful biological variation.

To further explore this complementarity, we examined the intersection of TVGs and SVGs with domain 10 differentially expressed (DE) genes and performed KEGG pathway enrichment (Fig. 6f). Both gene sets were enriched for immune-related functions, including chemokine, T cell receptor, and B cell receptor signaling, underscoring the immune-enriched nature of domain 10. However, TVGs showed stronger enrichment for pathways related to immune activation and inflammation, such as NF-kappa B signaling and primary immunodeficiency, suggesting ongoing immune responses. In contrast, SVGs were more enriched in pathways linked to immune regulation and structure, including PD-L1 checkpoint signaling and leukocyte migration, indicating spatially defined immune niches.

Overall, our results suggest that incorporating temporally variable gene signals provides complementary and biologically relevant insights to traditional spatial analyses. The proposed TSVG analysis, implemented through our method TESSA, integrates both pseudotime and spatial information, potentially enabling a more comprehensive characterization of complex tissue dynamics. While further validation is needed, our enrichment analysis suggests that TVGs contribute interpretable and non-redundant biological signals that may enhance understanding of spatial transcriptomic data.

#### 2.3.2 Human Lung Adenocarcinoma ST Data Analysis

We analyzed the 10x Visium spatial transcriptomics dataset from human lung adenocarcinoma (LUAD)[26]. Understanding the spatial architecture of cancer cells and the TME is crucial for lung cancer research. In this analysis, we observed that traditional SVGs, such as those identified by SPARK, tend to highlight large and strong spatial structures, such as tumor subclones, while potentially overlooking smaller but biologically significant regions, including tertiary lymphoid structures (TLS) and cancer-associated stromal areas. Accurate identification of both tumor subclones and normal tissue structures is essential for comprehensive cancer tissue analysis. We found that this balance is better achieved using uTSVGs identified by TESSA, which captures both large-scale tumor architecture and fine-grained features of the TME.

We used the lung adenocarcinoma in situ sample, TD8, as the main analysis sample, which contains 1,760 spots and 36,601 genes. After filtering out low-expressed genes, those expressed in fewer than 10% of spots, 11,138 genes remained. Pseudotime was inferred using Slingshot[11] based on cell type marker genes derived by scRNA-seq Human Lung Cell Atlas (HLCA)[27], resulting in three lineages that initiate from cancer regions (Fig. S6; see Supplementary File Pseudotime Estimation for details).

Using the TESSA Stage 1 overall test, we identified 4,955 significant genes, LOO removed 17 spurious correlated genes, leading to 4,938 uTSVGs at an FDR threshold of 0.05 by the Benjamini-Yekutieli correction. For comparison, SPARK identified 2,240 SVGs, with 2,166 overlapping with the uTSVGs. TESSA lineage-specific individual tests further identified 3,369 lineage 1 TVGs, 1,144 lineage 2 TVGs, 1,004 lineage 3 TVGs, and 757 SVGs (Fig. 7e).

**Fig. 7:**
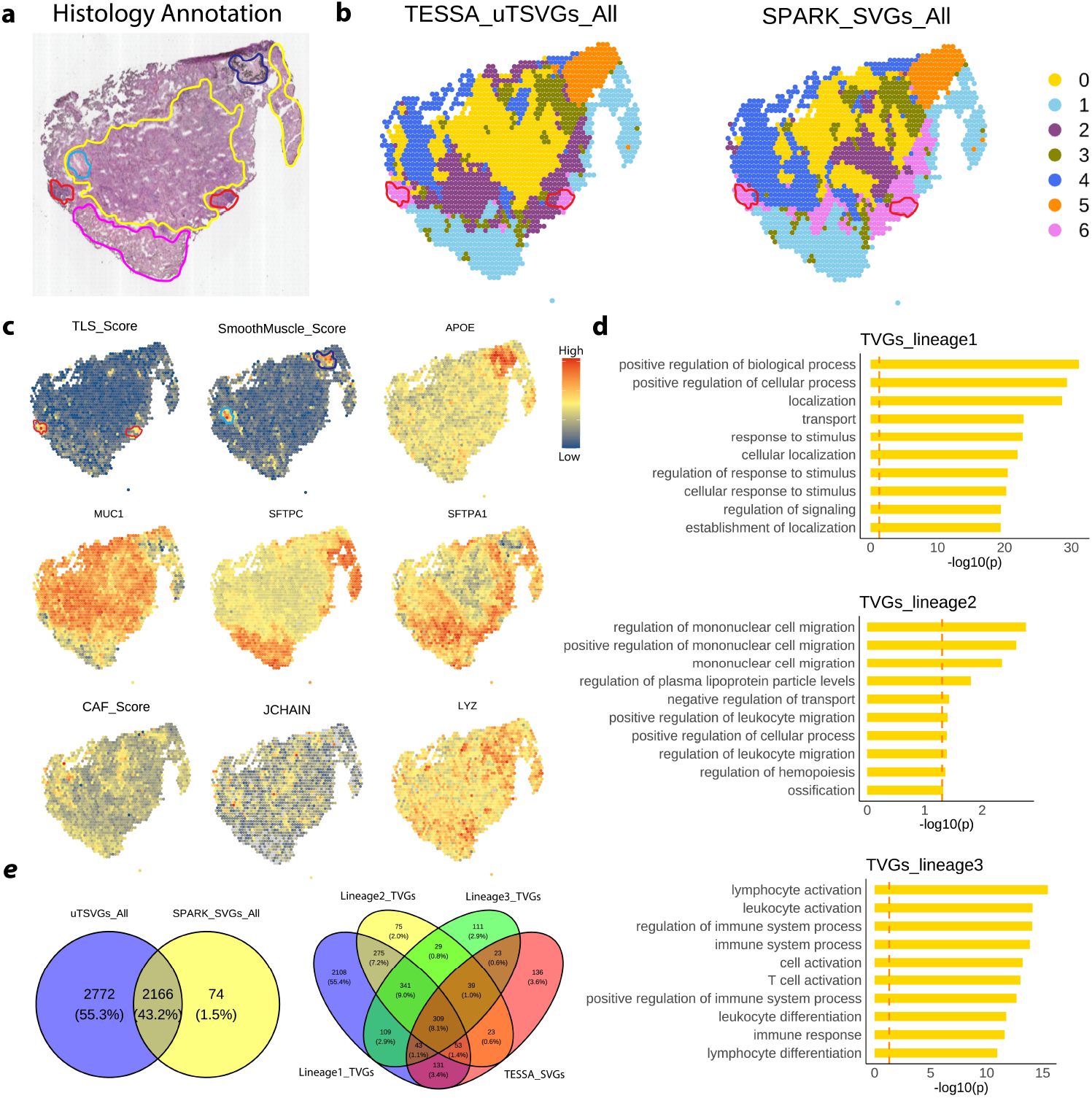
Results of Lung Adenocarcinoma Data. **a**. Histology domain annotation as ground truth. Yellow as cancer region, red as lymph region, pink as epithelial region, light blue and dark blue ones are stromal region. Light blue one is also identified as cancer vasculature region, and dark blue one is also identified as normal vasculature region. **b**. Domain detection results display the true and estimated domain annotations by SeuratPCA. Left: based on TESSA uTSVGs. Right: based on SPARK SVGs. **c**. Spatial pattern plots of log-normalized gene expression for cell type markers and biological structure signature scores(see details in supplementary files). **d**. GO BP term pathways enrichment of three lineage-specific unique TVGs. **e**. The left venn diagram shows the logical relationship between sets of genes identified by TESSA and SPARK. The right diagram shows the logical relationship between sets of lineage-specific TVGs and SVGs identified by TESSA stage 2 tests.

To evaluate whether incorporating temporally variable information improves downstream analysis, we compared domain detection performance using uTSVGs from TESSA and SVGs from SPARK. We applied both SeuratPCA [23] and SpatialPCA [4] with top 20 PCs, and used histological annotation (Fig. 7a) as reference. We focused on SeuratPCA domain result because it reveals more biological insights (see details of SpatialPCA domain results in Fig. S8). In general, the domain results based on TESSA uTSVGs and SPARK SVGs are similar and consistent with histological annotation, but domains based on TESSA uTSVGs outperform SPARK SVGs in some delicate but important structures.

##### uTSVG identified tertiary lymphoid structure (TLS) and tumor subclones

The domain results based on uTSVGs showed improved alignment with histological annotations, especially in small but biologically important tissue structures, compared to domains derived from SPARK SVGs (Fig 7.a-b). Moreover, TESSA uTSVGs reveal more information about tumor subclones (Fig 7.c).

Domains based on TESSA uTSVGs align well with histological annotations. For example, domain 6 successfully captured a small lymph node region, which was missed in the domain segmentation based on SPARK SVGs (Fig. 7b). Notably, domain 6 (violet) also showed high enrichment of TLS-score, suggesting it represents not only a lymph node but also a TLS (Fig. 7c), which is a known biomarker for immunotherapy response. The TLS score was calculated based on the expression of established TLS signature genes (*CETP, CCR7, SELL, LAMP3, CCL19, CCL21, CXCL9, CXCL10, CXCL11*, and *CXCL13*)[28, 29].

Domains based on TESSA uTSVGs reveal tumor subclones, aligned with cancer marker gene patterns. The expression pattern of cancer cell marker *MUC1* and normal epithelial marker *SFTPC* and *SFTPA1* both verify that domain 1 (skyblue) is normal epithelial[30], while domains 0,2,3,4 might be different cancer-related subclones. Domain 4 (blue) also shows high Cancer Associated Fibroblast (CAF) score and high expression for plasma cell marker gene *JCHAIN* [31], implying it could be a tumor stromal region with immune responses (score calculation in Supplementary File 3.3). Domain 2 (purple) shows high expression for *MUC1* and *SFTPA1* and low expression for *SFTPC*, indicating this is a transitional tumor zone, where Alveolar type 2 (AT2) derived tumor cells are de-differentiating or acquiring stem-like/progenitor properties. Myeloid marker *LYZ*[32] reveals that domain 3 (green) might be enriched in innate immune cells (Fig 7.a,b,c).

Although neither the domain results based on all TESSA uTSVGs nor those based on all SPARK SVGs fully recovered the stromal region annotated by the histologist, the use of the top 1,000 lineage-specific uTSVGs enabled detection of this stromal region (Fig. S6f). The discrepancy between data-driven domain segmentation and histological annotations should not simply be viewed as inaccuracy; rather, it reflects a fundamental challenge of spot-level spatial transcriptomics: the mixing of multiple cell types within each spot inherently limits cell-type-specific domain resolution. An example of this complexity is domain 5 (orange), which displays high expression of both smooth muscle-related genes and the tumor-associated macrophage (TAM) marker gene *APOE* [33, 34]. In this case, the TAM signal appears to dominate the identity of domain 5. In human lung cancer, *APOE* expression is significantly higher in tumor tissue compared to adjacent normal tissue and has been identified as a potential prognostic marker[33, 34]. Therefore, even if not included in histological annotations, TAM-enriched regions are biologically meaningful and important for LUAD analysis.

##### Lineage-specific TVGs are enriched in biologically distinct pathways

To assess the biological relevance of TVGs, we performed GO Biological Process enrichment analysis using lineage-specific unique TVGs, genes that are significantly variable over trajectory within a given lineage but not identified as SVGs (Fig. 7d). This allows us to focus on pseudotime-driven transcriptional programs specific to each lineage. In lineage 1, TVGs showed strong and broad enrichment of cell regulation and movement-related processes, suggesting a general cell type that is undergoing trafficking, secretory activity, or cellular remodeling. This lineage most likely represents stromal cells, like fibroblasts, or epithelial tumor cells undergoing cell secretory activity and structural functions[35, 36]. In lineage 2, the enriched pathways were more specific to immune migration-related pathways, especially the mononuclear cell reveals the cell trafficking related to innate immune myeloid cells, such as macrophages. Besides, plasma lipoprotein particle regulation is often linked to TAMs. Therefore, Lineage 2 represents a macrophage trajectory, indicating a pro-tumor or immune-modulatory infiltration process, typical in TAM biology[37]. In lineage 3, TVGs were strongly enriched for adaptive immune activation. Top enriched terms included T cell activation, lymphocyte differentiation, and positive regulation of immune system process, clearly pointing to lymphoid lineage cells, such as T cells and B cells. These patterns are consistent with the presence of TLS or immune infiltration at inflamed tumor boundaries, where adaptive immune responses play a critical role in tumor surveillance and elimination[38].

In summary, these results demonstrate that lineage-specific TVGs are not only statistically significant but also biologically meaningful. Different lineages show different patterns, ranging from supportive tissue, innate immune infiltration to adaptive immune activation. This variety demonstrates the ability of TESSA to uncover pseudotemporal trajectories that are relevant to tissue organization and immune-tumor dynamics. By focusing on TVGs, we reveal dynamic processes that are not fully captured by spatial-only analyses, further reinforcing the biological meaningfulness and utility of TVGs in characterizing complex tissue states.

#### 2.3.3 Multiple Sample Integration

TESSA enables multi-sample integration to identify uTSVGs. We applied TESSA on two published datasets obtained using different spatial transcriptomics technologies to illustrate the benefit of TESSA that contributes to multi-sample uTSVGs detection, which is shown to improve domain detection accuracy. The two datasets include the STARmap single cellular resolution ST dataset from the mouse primary visual cortex[39] and the 10x Visium spot-level ST dataset from the six-layered human dorsolateral prefrontal cortex[40].

##### uTSVGs improve domain detection accuracy for the mouse primary visual cortex data

We examined the mouse STARmap primary visual cortex data[39]. The cells on all tissue sections were carefully annotated into four distinct layer structures that included L1, L2/3, L5, and L6. We used this dataset to illustrate the benefit of TESSA that contributes to multi-sample uTSVG detection. The gene expression count data are measured across 3 samples, BZ5(1049 cells), BZ9(1053 cells) and BZ14(1088 cells), with a common gene panel of 166 genes. We used Slingshot[11] to estimate the pseudotime inspired by 15 signature genes for cell types found in the mouse primary visual cortex. We removed low-expressed genes that do not express at more than 10% of spots, resulting in 131 genes. There are 12 signature genes in the remaining gene set (details in Supplementary File Pseudotime Estimation).

The TESSA overall test identified 111 uTSVGs with p-values adjusted by the Benjamini-Yekutieli correction at the 0.05 FDR level. The LOO strategy removed one signature gene *Pax6* from the significant gene list, resulting in 110 significant uTSVGs. SPARK identified 63 SVGs, all of which are also identified by the TESSA overall test. To evaluate whether incorporating temporally variable information across multiple samples can improve downstream domain analysis, we compared domain detection accuracy using uTSVGs identified by TESSA, as well as SVGs detected by SPARK, as input features. The multiple sample extension of SpatialPCA[4] was applied as the main domain detection algorithm. The adjusted Rand index (ARI)[41] and Normalized Mutual Information (NMI) were used for evaluation. We achieved an ARI of 0.62 and NMI of 0.59 with uTSVGs identified by TESSA for all three samples, while SPARK SVGs yielded lower accuracy with an ARI of 0.49 and NMI of 0.50 (Fig. 8d). For example, in all three samples, using SPARK SVGs as input, it falsely classified more L1, L2/L3 layer cells into L4 layer. In sample BZ14, SPARK SVGs identified many L1 and L4 layer cells, which are supposed to be L5 layer cells. In contrast, TESSA uTSVGs resulted in a cleaner L5 domain (Fig. 8a,b,c).

**Fig. 8:**
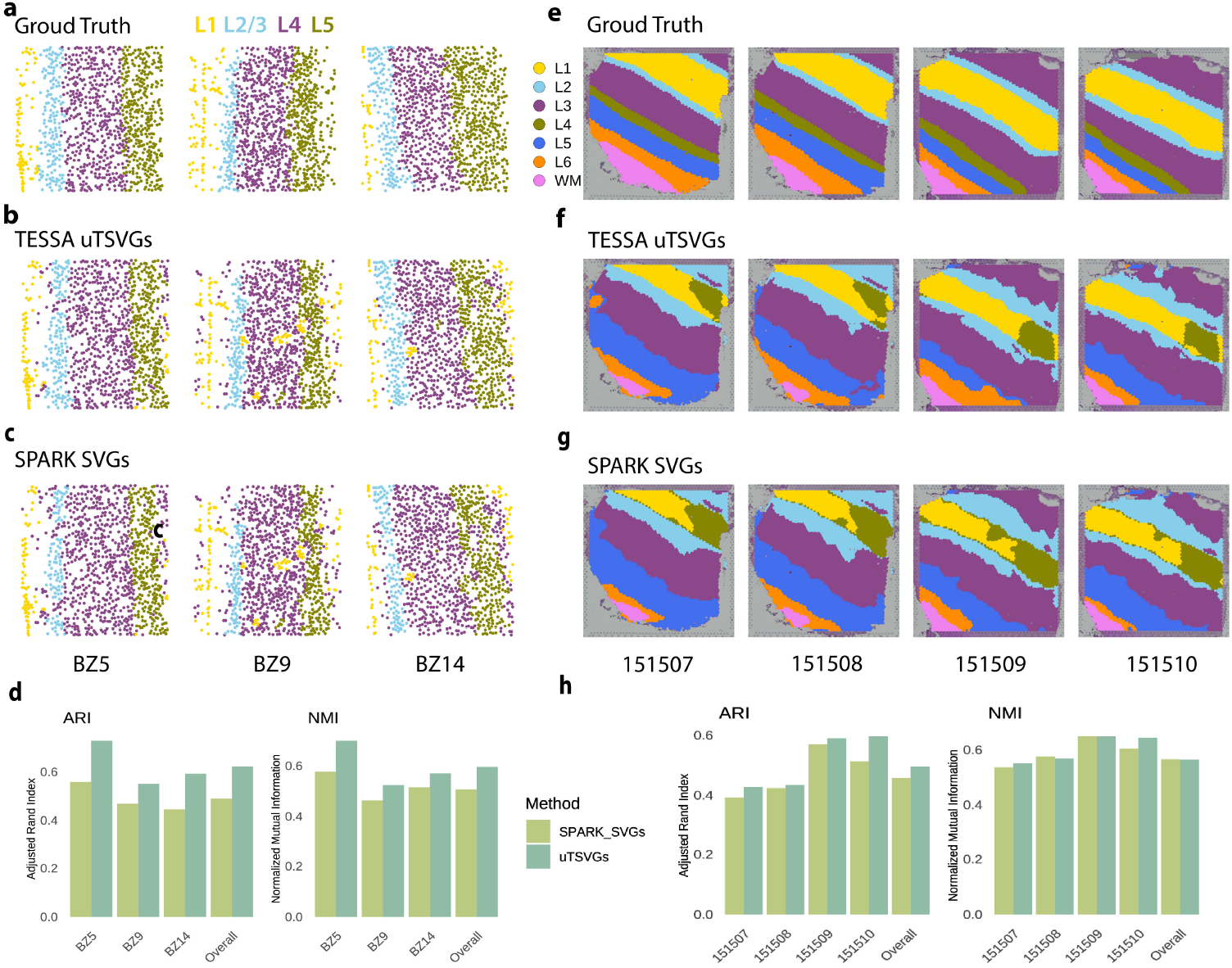
Results of Mouse Cortex Data and Human Cortex Data. **a**. Ground truth domain annotation for mouse cortex data. **b**. Domain detection results based on TESSA TSVGs. **c**. Domain detection results based on SPARK SVGs. **d**. Barplots of ARI and NMI show the accuracy of domain detection for sample BZ5, BZ9, BZ15 and all samples together (Overall). Higher ARI and NMI mean more accurate domain detection. **e**. Ground truth domain annotation for human cortex data. **f**. Domain detection results based on TESSA TSVGs. **g**. Domain detection results based on SPARK SVGs. **h**. Barplots of ARI and NMI show the accuracy of domain detection for individual samples and all samples together (Overall).

##### uTSVGs improve domain detection accuracy for human dorsolateral prefrontal cortex data

We also examined human dorsolateral prefrontal cortex data[40]. The tissue sections are annotated as seven spatial domains, including six cortical layers (L1-L6) and white matter. We used this dataset to illustrate that TESSA uTSVGs across samples improved domain detection accuracy compared to the results by SPARK SVGs.

We used 4 samples, namely 151507 (4,226 spots), 151508 (4,384 spots), 151509 (4,789 spots), and 151510 (4,634 spots), for the analysis, which contain 18,033 spots and 33,538 genes in total. After filtering out low-expressed genes, those expressed in fewer than 10% of spots, 3,783 genes remained. Pseudotime was inferred using Slingshot[11] based on 28 signature genes (see Supplementary File 3.1).

The TESSA overall test identified 1,521 uTSVGs with p-values adjusted by the Benjamini-Yekutieli method to meet an FDR level of 0.05. After implementing the LOO strategy, three signature genes were removed due to potential double dipping, resulting in 1,519 uTSVGs. SPARK identified 685 SVGs, of which 682 genes are also identified by the TESSA overall test. Again, we applied the multiple sample extension of SpatialPCA[4] for domain detection. We achieved an ARI of 0.49 and NMI of 0.56 with uTSVGs identified by TESSA for all four samples together, while SPARK SVGs yielded slightly lower accuracy with ARI of 0.45 and NMI of 0.56 (Fig. 8h). Compared to domain results based on SPARK SVGs, uTSVGs identified L2 layer(skyblue) better in samples 151509 and 151510 and shrunk the size of artificial green cluster in L1 layer (gold) (Fig. 8e,f,g).

## 3 Discussion

In this work, we introduced TESSA, a unified statistical framework developed to detect temporally and spatially variable genes (TSVGs) in spatial transcriptomics. By integrating gene expression, spatial location, and pseudotime through a linear mixed-effect model, TESSA enables the identification of uTSVGs in an initial stage, followed by a second stage dedicated to further distinguishing TVGs and SVGs. Its design ensures robustness in various data generation scenarios and maintains invariance to spatial coordinate rotation[42]. In addition, the proposed LOO strategy effectively addresses the well-known double-dipping issue in pseudotime-related inference and provides a promising solution for broader applications.

TESSA extends pseudotime-based differential expression testing to spatial transcriptomics analysis by leveraging user-defined pseudotime. It is important to note that several methods have been developed to infer spatial pseudotime, which integrates both gene expression and spatial coordinates, thereby adapting traditional pseudo-time inference to ST data. Notable examples include SpaceFlow[43], PSTS[44], and PearlST[45]. In this study, we opted to estimate pseudotime using the scRNA-seq-based method Slingshot[11], applied to user-defined signature genes. This decision was based on two considerations: first, we observed that spatial pseudotime methods tend to over-smooth anatomical structures particularly in tumor tissues (Fig. S4); second, spatial pseudotime may introduce confounding effects in the detection of SVGs. While we employed Slingshot in this work, we emphasize that the choice of pseudo-time estimation method is not central to our framework, and users are free to integrate alternative approaches as needed.

Additionally, utilizing uTSVGs identified by TESSA enhances spatial domain detection on both single and multiple samples, compared to SVGs detected by other methods such as SPARK, as demonstrated in the analysis of the real data. The down-stream functional enrichment analysis further underscores the advantages of using TESSA in capturing functionally relevant genes that vary along dynamic trajectories.

Despite its strengths, TESSA has areas for improvement. Firstly, we applied Gaussian kernels on both spatial and temporal covariance matrices, with a common bandwidth selected by a rule-of-thumb criterion[46]. Kernel and bandwidth selection maintains a tricky question; for further improvement, we can also apply the Cauchy p-value combination test[47] to aggregate the power for multiple kernels and band-widths to increase detection power. Secondly, TESSA is designed for single lineage. In theory, we are able to include multiple *L* lineages in one model as additional variance components, such as 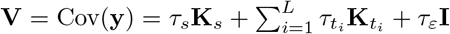. But this will increase computational complexity and require another comprehensive investigation, which will be considered in our future work. Thirdly, TESSA incorporates matrix decomposition and inversion on the spatial and temporal kernel matrix; these kernel operations could be slow as spot size increases. In our future work, we will explore kernel approximation techniques such as the nearest neighbor idea implemented in nnSVG [48], to improve scalability.

## 4 Methods

### 4.1 The Model

Our goal is to identify genes that exhibit temporally and spatially variable expression patterns, referred to as TSVGs, and to further assess whether these genes show temporal or spatial variation. Suppose we have spatial transcriptomics expression data for *p* genes measured across *n* spots (or cells) of a 2-D tissue. Let the spatial coordinates be denoted by **s** = (*s*_*i*1_, *s*_*i*2_)_*n×*2_, and the corresponding pseudotime values by **t** = (*t*_*i*_)_*n×*1_, *i* = 1, … , *n*. Let 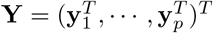 denotes the normalized gene expression matrix, where **y**_*j*_ = (*y*_*j*1_, … , *y*_*jn*_)^*T*^ represents the normalized expression of gene *j* across all spots (or cells). Here, *y*_*ji*_ denotes the normalized expression level of gene *j* in spot (or cell) *i*.

For the *p* genes, we divide them into two groups: ℳ denotes a user-defined set of *m* signature genes for pseudotime estimation, and 𝒦 denotes the remaining *k* = *p* − *m* genes. In this work, we infer pseudotime using the Slingshot[11] algorithm, based solely on the expression profiles of genes in ℳ. Users can use other pseudotime estimation algorithms.

For notation simplicity, when referring to an individual gene, we omit the subscript index *j* and denote its normalized gene expression as **y** = (*y*_1_, …, *y*_*n*_)^*T*^ . Based on this setup, we establish a variance component model to explicitly decompose the contribution of each gene’s expression into components associated with spatial and temporal variability, i.e.,

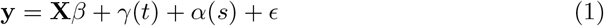

where **X** denotes a covariate matrix with *v* dimensions, *γ*(*t*) ∼ *MV N* (0, *τ*_*t*_**K**_*t*_), *α*(*s*) ∼ *MV N* (0, *τ*_*s*_**K**_*s*_), **K**_*t*_ and **K**_*s*_ are temporal and spatial kernel matrcies cal-culated based on the estimated pseudotime and spatial locations, respectively, and *ϵ* ∼ *MV N* (0, *σ*^2^**I**). Thus, we have **y** ∼ *MV N* (**X*β*, V**), where **V** = Cov(**y**) = *τ*_*s*_**K**_*s*_ + *τ*_*t*_**K**_*t*_ + *τ*_*ε*_**I**, and (***β***, *τ*_*s*_, *τ*_*t*_, *τ*_*ε*_) are the unknown parameters to estimate. The remaining parameters (e.g., kernel bandwidth) define covariance structures that capture different types of spot-level similarity. The key intuition is that a gene that exhibits local spatial correlation is captured by the spatial kernel **K**_*s*_ and a gene associated with pseudotime progression is captured by the temporal kernel **K**_*t*_. **K**_*s*_ and **K**_*t*_ are pre-defined kernel matrices, while *τ*_*s*_ and *τ*_*t*_ are variance components. The details about kernel construction can be found in Supplementary File 1.4.

The model aims to dissect gene expression variance into components driven by spatial and temporal effects, thereby offering interpretable insights for downstream biological analyses. The detailed estimation steps can be found in Supplementary File 1.1.

### 4.2 Hypothesis Testing

We first conduct the stage 1 overall test to assess if a gene shows any temporal or spatial variation with the following hypotheses:

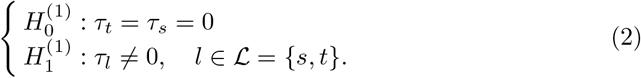

Under the null hypothesis, the model simplifies to **y** ∼ *MV N* (**X*β***, *σ*^2^**I**), which is a linear model with only fixed effects and an error term, indicating that the gene expression does not show either spatial or temporal variability. If the null hypothesis is rejected, we call the gene a uTSVG, which could be an SVG, TVG or both. A score statistic is constructed for detecting uTSVGs, and the corresponding p-value can be computed through a scaled chi-square distribution. We examine one gene at a time, and once the p-values for all genes are obtained, a false discovery rate (FDR) control procedure (e.g., the Benjamini-Yekutieli procedure) is performed across all genes to declare the final significant gene list.

If the null 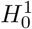 is rejected, we proceed to the stage 2 test with the following hypotheses:

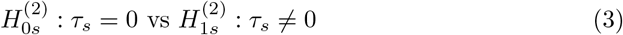

to test if a gene is an SVG, and

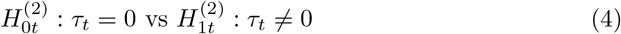

to test if a gene is a TVG. Similarly, a score statistic is constructed to detect TVGs and SVGs, and the associated p-value can be computed through approximation by a scaled chi-square distribution. The details about the testing procedures can be found in Supplementary File 1.2 and 1.3.

### 4.3 Double Dipping Correction

In real-world data, the oracle pseudotime is unknown. If a gene is used for pseudotime estimation, then tested for temporal variation based on the estimated pseudotime, this will cause an inflated type I error due to spurious correlation. This is a problem known as double dipping, which is inevitable when identifying cluster or pseudotime-associated differentially expressed genes[17].

To address this, we propose the LOO strategy, designed to alleviate the double-dipping issue in both Stage 1 (uTSVG detection) and Stage 2 (TVG detection) testing. Unlike commonly used data-splitting approaches, such as cross-validation to split data samples[49], LOO excludes the test gene from the pseudotime estimation step. This design ensures that no spurious correlation is introduced between the estimated pseudotime and the test gene, even in the absence of a true underlying trajectory. Moreover, because double dipping arises from reusing gene information, LOO only needs to be applied to the subset of *m* user-defined signature genes used to infer pseudotime. This focused application balances statistical rigor with computational efficiency. The full procedure of Stage 1 test with LOO implementation is summarized in Algorithm 1. In this strategy, we assume no single signature gene dominates the pseudotime estimation. In practice, the number of signature genes is usually in the tens or hundreds. The impact of pseudotime estimation by removing one gene is usually minimal. Our simulation results demonstrate that the LOO strategy not only alleviates inflated type I error but also improves testing accuracy (see Fig.4. a,b; Fig.5. a,b,e,f).

#### Algorithm 1

Stage 1 Hypothesis Testing with LOO

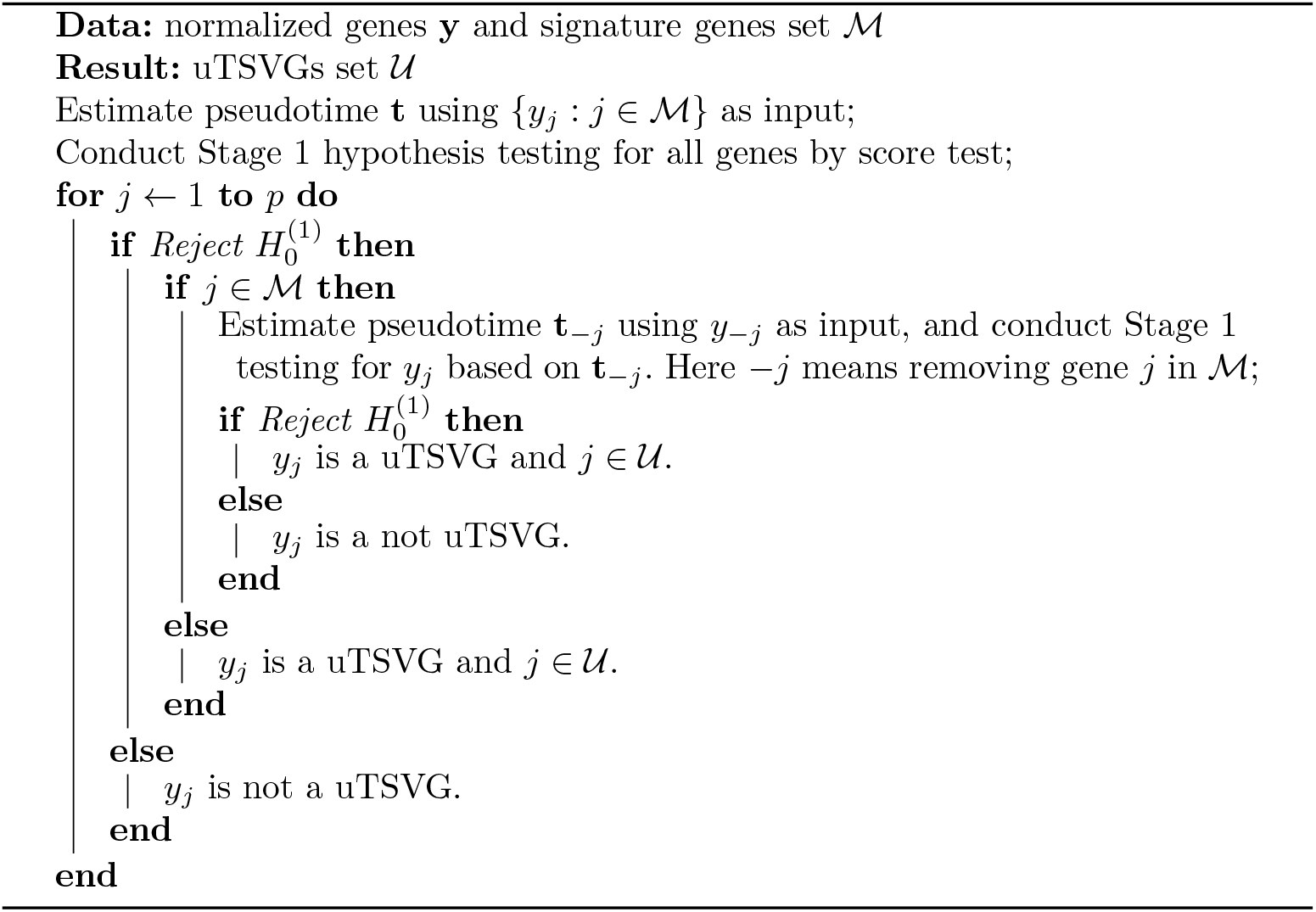

### 4.4 Extension to Multiple Sample Integration

Multiple samples are increasingly generated due to the decreasing cost of ST technologies. Our TESSA framework can be extended to integrate multiple tissue samples for TSVG detection. Suppose we have *Q* tissue samples, and 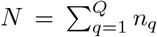 spots, where *n*_*q*_ denotes the number of spots in each sample. In the extension, we first regress out sample batch effect in the variance-stabilizing transformation[21] step, to correct sample mean difference. Assuming samples are independent, we construct the spatial kernel matrix as a block diagonal matrix 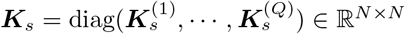 based on the location information of each tissue sample. As for the construction of the temporal kernel matrix, unlike spatial locations, the temporal information of spots/cells in different samples may not be independent. For example, normal epithelial cells in different samples may share similar temporal information and should not be treated as independent. Therefore, considering pseudotime is an abstract distance of gene expression similarity, we use the pseudotime information of all spots to construct the large temporal kernel matrix ***K***_*t*_ ∈ ℝ^*N ×N*^ . The constructed ***K***_*s*_ and ***K***_*t*_ are then fitted to model (1) for further parameter estimation and testing following the same procedure laid out in previous sections. As illustrated in our real data application, TESSA can be applied to both the single-cell resolution (e.g., the Starmap mouse cortex data with three samples) and spot-level data (e.g., the human dorsolateral prefrontal cortex data with four samples), assuming the sample size is not too large.

### 4.5 Comparison of Methods

We compared TESSA with the Gaussian version of SPARK (SPARK-G)[21] for uTSVGs detection, and with Countsplit[17] and DataThin[18] for double dipping correction. Both TESSA and SPARK-G are designed for normalized gene expression data. For notation simplicity, SPARK is used in this work to replace SPARK-G. To ensure consistency in preprocessing, we modified the gene and cell filtering criteria in SPARK to match TESSA gene sets. Countsplit is designed for count data, while DataThin extends this framework to normalized continuous data. Therefore, for the Negative Binomial simulation scenario and real datasets, we applied Countsplit to split the raw count expression data into training and testing set. For the Normal simulation scenario, we applied DataThin to split the continuous expression data within each spot. In both cases, pseudotime was estimated using the training data, and hypothesis testing was conducted on the test data.

Note that this work is not aimed at performing an exhaustive benchmark against all existing SVG detection methods. We selected SPARK as a representative comparison because it is a popular approach widely adopted in many applications for SVG detection. Like our approach, SPARK is based on a variance component framework; however, it constructs spatial kernels using a combination of five Gaussian and five cosine kernels with varying bandwidths and employs the Cauchy combination test for p-value aggregation to boost statistical power. In our analysis, TESSA leverages a simpler kernel design, using a single spatial Gaussian kernel and a temporal Gaussian kernel with a fixed bandwidth selected by the rule-of-thumb criterion[46, 50]. This comparison is intended to specifically highlight the contribution and importance of incorporating temporal information in detecting spatiotemporally variable genes. Practically, we can also borrow the SPARK idea to incorporate multiple bandwidth and use the Cauchy p-value combination to enhance statistical power.

We selected Countsplit and DataThin as benchmarks for double dipping correction, because they clearly formulated the double dipping issue into a statistical framework and are easy to implement. PseudotimeDE[16] has been developed for pseudotime DE gene analysis for scRNA-seq data without spatial information. We did not compare with it for two reasons: 1) it could yield anti-conservative p-values in the absence of a true trajectory[17], and 2) it is computationally intensive, requiring hundreds to thousands of permutations to estimate the null distribution. Countsplit applies Bernoulli sampling to split the gene expression data into two independent training and testing matrices, such that their sum recovers the original matrix. Pseudotime is estimated from the training matrix, and gene testing is performed on the testing matrix. Under the Poisson assumption, the two matrices are independent; thus, Countsplit avoids spurious correlation and controls type I error as shown in the paper[17]. DataThinning [18] generalizes this approach to other convolutional closed distributions, such as the Normal distribution.

## 5 Simulation Design

We designed two data generation scenarios to evaluate the robustness of TESSA on data generated from a Negative Binomial distribution and a Normal distribution. The first scenario uses data generated from a Negative Binomial distribution for count data, while the second uses a Normal distribution to simulate normalized expression data. The Normal scenario serves as an idealized setting to verify the efficacy of TESSA, whereas the Negative Binomial scenario provides a more realistic benchmark for assessing type I error control and statistical power as in real data applications.

### 5.1 Data Generation: Normal Scenario

We designed the Normal scenario to evaluate the performance of TESSA under an idealized setting. To generate realistic spatial and temporal structures, we referred to human lung cancer spatial transcriptomics data. Specifically, we extracted the estimated pseudotime from a real lineage trajectory in the LUAD data (see pseudotime details in section 2.3.2 and Supplementary File 3.1) and treated it as the oracle pseudotime. Together with the spatial coordinates of each spot, these were used as inputs to generate simulated gene expression values.

The expression of a gene of interest, denoted by ***y***, was simulated from a multivariate normal distribution:

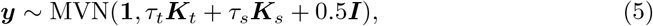

The parameters *τ*_*t*_ and *τ*_*s*_ control the strength of the temporal and spatial signals, and the residual variance is fixed at 0.5. To assess the false positive control, we set *τ*_*t*_ = *τ*_*s*_ = 0 to simulate null genes with no spatial and temporal effect. To evaluate testing power, we varied the signal strength by setting *τ* ∈ {0.3, 0.5}, and decomposed the signal into spatial and temporal components: *τ*_*s*_ = *ατ* , *τ*_*t*_ = (1 − *α*)*τ* . This allowed us to generate SVGs (*α* = 1), TVGs (*α* = 0), TSVGs (e.g., *α* = 0.5), and null genes (*τ* = 0). For each type of expression pattern, we simulated 50 genes and repeated the procedure 20 times, resulting in 1,000 gene replicates in each case. Pseudotime was estimated using the Slingshot algorithm (Fig. S1a,b), and we selected 20% of the genes as user-defined signature genes for pseudotime inference. This set consisted of randomly selected 80% TVGs, 10% TSVGs, and 10% null genes.

### 5.2 Data Generation: Negative Binomial Scenario

Motivated by the tumor progression trajectory observed in biological applications, we constructed a spatial pattern with four domains: D0 (normal tissue), D1 (tumor in situ), D2 (invasive tumor), and D3 (an unrelated domain independent of tumor progression) (Fig. 9a). Spatial coordinates were sampled from the unit square [0, 1]^2^, and the oracle pseudotime was drawn from the range [0, 1]. We simulated the data such that pseudotime similarity are not necessarily related to spatial location similarity. That is to say, even if spots are spatially far from each other, pseudotime values could be similar as long as they are within the same developmental domain, such as in D1. For each domain, pseudotime values were generated using truncated Normal distributions (Fig. 9b) with values 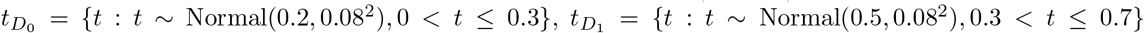 and 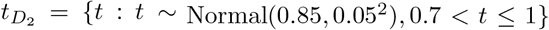. We could also simulate oracle pseudotime using a uniform distribution. However, pseudotime distributions are typically more complex, with certain regions being densely populated while others are sparse. Therefore, we did not use a uniform distribution for this purpose (Fig. S5).

**Fig. 9:**
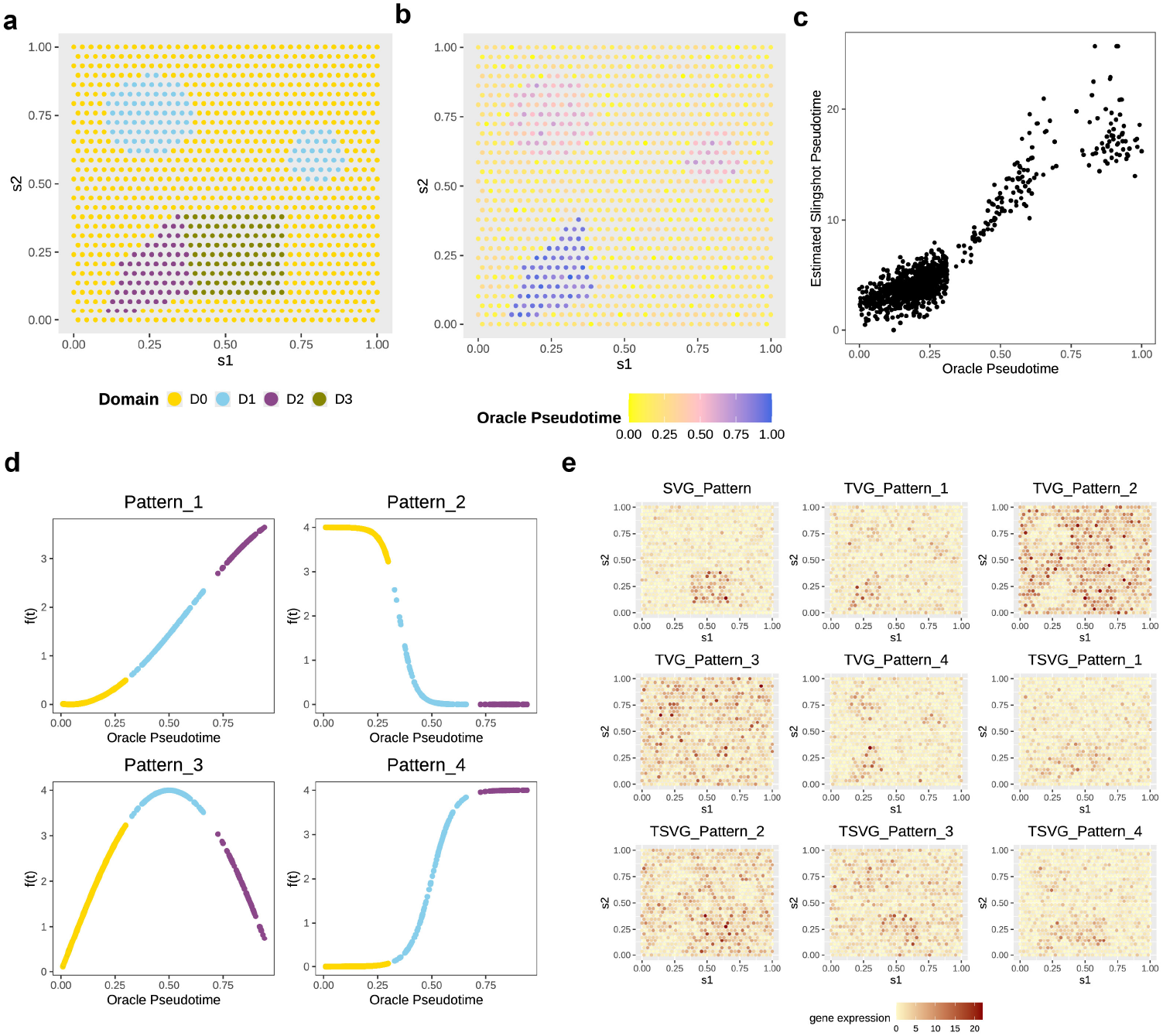
Simulation Data Generation under the Negative Binomial Scenario. Visualization of spatial domains. Domains D0, D1, and D2 are associated with temporal effects, while domain D3 exhibits a spatial effect without a temporal component. Spatial distribution of the oracle pseudotime. **c**. Scatter plot comparing the oracle pseudotime and the pseudotime estimated by Slingshot. **d**. The four predefined oracle temporal patterns of ***f* (*t*). e**. Spatial raw count expression patterns of representative genes corresponding to the nine signal patterns.

We assume the raw count expression of a gene, denoted as 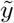, follows a Negative Binomial (NB) distribution,

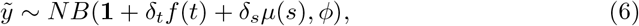

where *ϕ* is the dispersion parameter, *δ*_*s*_ represents the spatial signal strength and *δ*_*t*_ denotes the temporal signal strength. The function *f* (*t*) encodes the temporal pattern associated with pseudotime *t*, and *μ*(*s*_*i*_) = 𝕀{*s*_*i*_ ∈ *D*3} is an indicator function denoting whether spot *s*_*i*_ belongs to the spatially variable domain D3. We designed four distinct temporal patterns for *f* (*t*) to assess the method’s power under varying temporally dynamic expression trends (see Fig. 9d).

In this parameterization of the Negative Binomial distribution, the mean is given by *μ*_*NB*_ = 1 + *δ*_*t*_*f* (*t*) + *δ*_*s*_*μ*(*s*) and the variance is given by 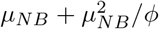. This relationship shows that the variance is directly influenced by the value of the dispersion parameter *ϕ*. If 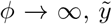 follows a Poisson distribution. We developed several simulation scenarios to assess the type I error control and testing power under different gene expression patterns and gene expression dispersions (i.e., 0.7, 1, 1.5, with higher dispersion corresponding to lower variation).

To assess false positive control, we set *δ*_*t*_ = *δ*_*s*_ = 0. To evaluate power, we used *δ* ∈ {2, 3} as the overall signal strength and decomposed the overall signal strength into spatial and temporal components: *δ*_*s*_ = *αδ* and *δ*_*t*_ = (1 − *α*)*δ*, where *α* controls the proportions of TVG and SVG distribution. This allowed us to simulate three types of signal genes: SVGs (*α* = 1), TVGs (*α* = 0), and TSVGs (*α* = 0.5), as well as noise genes (*δ* = 0). When *α* = 1, we simulate SVGs as *δ*_*t*_ = 0 and *δ*_*s*_ = 1, with *μ*_*NB*_ = 1 + *δ*_*s*_*μ*(*s*). In this case, *δ*_*s*_ serves as the fold change in the SVG simulation of SPARK[21]. We simulated 50 genes for each type of expression pattern (Fig. 9e), and repeated this procedure 20 times, resulting in 1,000 gene replicates for each of the 9 TSVG types. Then we applied a variance-stabilizing transformation [21] to convert raw count data into approximately Normal-distributed expression values. Pseudotime was estimated using the Slingshot algorithm(Fig. 9c), and we selected 20% of the genes as user-defined signature genes for pseudotime inference. This set consisted of randomly selected 80% TVGs, 10% TSVGs, and 10% noise genes.

### 5.3 Simulation 1: Evaluation of the Overall Test

The first set of simulations evaluated the performance of the Stage 1 overall test for TSVGs detection. As TESSA relies on pseudotime information to detect TSVGs, we tested TESSA using both true pseudotime (TESSA-oracle) and estimated pseudotime by slingshot (TESSA-slingshot). We evaluated the false positive control of TESSA with the LOO strategy (TESSA-slingshot-LOO) and Countsplit (TESSA-slingshot-countsplit).

For the null case, we set *τ* = 0 (for the Normal scenario) or *δ* = 0 (for the NB scenario) to simulate gene expression in the absence of spatial and temporal signals. Type I error control was evaluated under three different dispersion parameter values. In each of the 20 replicates per dispersion setting, we simulated 500 noise genes and randomly selected 40 of them as signature genes for pseudotime estimation. This resulted in a total of 10,000 noise gene replicates per dispersion level, repeated across all three dispersion settings.

For the alternative case, we set *δ* = 2 or *δ* = 3 in the NB scenario and *τ* = 0.3 or *τ* = 0.5 in the Normal scenario, to simulate gene expressions with spatial and temporal signals, allowing us to evaluate statistical power and TSVG detection accuracy. For each *τ* value in the Normal scenario and each combination of *δ* and dispersion in the NB scenario, we conducted 20 replicates. In each replicate, we simulated 50 genes for each of the 9 signal patterns in the NB scenario (and each of the 3 signal patterns in the Normal scenario), along with 50 noise genes. Among these, 40 genes were randomly selected as signature genes for pseudotime estimation, comprising 32 TVGs, 4 TSVGs, and 4 noise genes. This resulted in a total of 3,000 uTSVGs and 1,000 noise genes for each *τ* value in the Normal scenario and a total of 9,000 uTSVGs and 1,000 noise genes for each *δ* and dispersion in the NB scenario.

### 5.4 Simulation 2: Evaluation of the Individual Test

This simulation was designed to evaluate the performance of the Stage 2 individual test for detecting TVGs and SVGs, to assess the model performance for both type I error control and statistical power.

We used the same data generation mechanism as in Simulation 1 for both the null and alternative cases. Since TVG detection in TESSA relies on pseudotime information, we evaluated type I error control, detection accuracy, and power under different methods: TESSA-oracle, TESSA-slingshot, TESSA-slingshot-countsplit, and TESSA-slingshot-LOO. Since SVG detection does not involve pseudotime and is not affected by double dipping, we evaluated SVG detection performance only under TESSA-oracle and TESSA-slingshot.

## Data availability

The human pancreatic cancer data[22] can be found through dbGaP with accession number PRJNA1124001 and the image files are available at https://zenodo.org/records/13379726. The human lung cancer data[26] is deposited at Gene Expression Omnibus (GEO) under the accession codes GSE189487. The mouse primary visual cortex data[39] can be found at https://github.com/zhengli09/BASS-Analysis/tree/master/data. The human dorsolateral prefrontal cortex data [40] can be found at https://github.com/LieberInstitute/HumanPilot.

## Code availability

The R code used to develop the model, perform the analyses and generate results in this study is publicly available on GitHub at https://github.com/Cui-STT-Lab/TESSA under GPL-3.0 license.

## Acknowledgement

This study was supported by the High-Performance Computing Center (HPCC) at Michigan State University.

## Author contribution

Y.C. conceived the idea. Y.W. and Y.C. designed the experiments. Y.W. developed the algorithm, implemented the software, performed simulations, and analyzed real data. H.S. helped with algorithm development. Y.X. and N.S. helped with data interpretation. Y.W. and Y.C. wrote the manuscript.

## Competing interests

The authors declare no competing interests.

## Notes

### Competing Interest Statement

The authors have declared no competing interest.

