## Supplemental File for "TESSA: A unified model to detect trajectory-preserved and spatially-variable genes in spatial transcriptomics"

**4 Supplementary Figures** **11**

### 1 Details of TESSA

#### 1.1 The Estimation

Denote $\boldsymbol{\tau} = (\tau_s, \tau_t, \tau_\varepsilon)^T$ and $\boldsymbol{\theta} = (\boldsymbol{\beta}^T, \boldsymbol{\tau}^T)^T$, then the likelihood function for $\boldsymbol{\theta}$ is

$$L(\boldsymbol{\theta}; \mathbf{y}) = (2\pi)^{-\frac{n}{2}} |\mathbf{V}|^{-\frac{1}{2}} \exp\{-\frac{1}{2}(\mathbf{y} - \mathbf{X}\boldsymbol{\beta})^T \mathbf{V}^{-1} (\mathbf{y} - \mathbf{X}\boldsymbol{\beta})\}. \tag{1}$$

Then, the log-likelihood function for $\boldsymbol{\theta}$ is given by

$$\ell(\boldsymbol{\theta}; \mathbf{y}) = -\frac{n}{2}\log(2\pi) - \frac{1}{2}\log|\mathbf{V}| - \frac{1}{2}(\mathbf{y} - \mathbf{X}\boldsymbol{\beta})^T \mathbf{V}^{-1}(\mathbf{y} - \mathbf{X}\boldsymbol{\beta}) \tag{2}$$

Take the first derivative of the log likelihood function with respect to $\boldsymbol{\beta}$ and set it to 0, we obtain the restricted maximum likelihood (REML) estimator as

$$\hat{\boldsymbol{\beta}} = (\mathbf{X}^T \mathbf{V}^{-1} \mathbf{X})^{-1} \mathbf{X}^T \mathbf{V}^{-1} \mathbf{y}. \tag{3}$$

Profiling out $\boldsymbol{\beta}$ and taking logarithm gives us the restricted log likelihood function for $\boldsymbol{\tau}$,

$$\begin{aligned} \ell^R(\boldsymbol{\tau}; \mathbf{y}) &\propto -\frac{1}{2}\log|\mathbf{V}| - \frac{1}{2}\log|\mathbf{X}^T\mathbf{V}^{-1}\mathbf{X}| - \frac{1}{2}(\mathbf{y} - \mathbf{X}\hat{\boldsymbol{\beta}})^T\mathbf{V}^{-1}(\mathbf{y} - \mathbf{X}\hat{\boldsymbol{\beta}}) \\ &\propto -\frac{1}{2}\log|\mathbf{V}| - \frac{1}{2}\log|\mathbf{X}^T\mathbf{V}^{-1}\mathbf{X}| - \frac{1}{2}\mathbf{y}^T\mathbf{P}\mathbf{y}, \end{aligned} \tag{4}$$

where $\mathbf{P} = \mathbf{V}^{-1} - \mathbf{V}^{-1}\mathbf{X}(\mathbf{X}^T\mathbf{V}^{-1}\mathbf{X})^{-1}\mathbf{X}^T\mathbf{V}^{-1}$.

Then, the score vector of $\boldsymbol{\tau}$ is given by

$$\mathbf{S}(\boldsymbol{\tau}; \mathbf{y}) = \big(S(\tau_s; \mathbf{y}), S(\tau_t; \mathbf{y}), S(\tau_\epsilon; \mathbf{y})\big)^T, \tag{5}$$

where for $l \in \mathcal{L} = \{s, t\}$,

$$\begin{aligned} S(\tau_l; \mathbf{y}) &= \frac{\partial \ell^R(\boldsymbol{\tau}; \mathbf{y})}{\partial \tau_l} \\ &= -\frac{1}{2}\mathrm{tr}(\mathbf{V}^{-1}\mathbf{K}_l) + \frac{1}{2}\mathrm{tr}\big[(\mathbf{X}^T\mathbf{V}^{-1}\mathbf{X})^{-1}\mathbf{X}^T\mathbf{V}^{-1}\mathbf{K}_l\mathbf{V}^{-1}\mathbf{X}\big] + \frac{1}{2}\mathbf{y}'\mathbf{P}\mathbf{K}_l\mathbf{P}\mathbf{y} \\ &= -\frac{1}{2}\mathrm{tr}\Big\{\big[\mathbf{V}^{-1} - \mathbf{V}^{-1}\mathbf{X}(\mathbf{X}^T\mathbf{V}^{-1}\mathbf{X})^{-1}\mathbf{X}^T\mathbf{V}^{-1}\big]\mathbf{K}_l\Big\} + \frac{1}{2}\mathbf{y}'\mathbf{P}\mathbf{K}_l\mathbf{P}\mathbf{y} \\ &= -\frac{1}{2}\mathrm{tr}(\mathbf{P}\mathbf{K}_l) + \frac{1}{2}\mathbf{y}'\mathbf{P}\mathbf{K}_l\mathbf{P}\mathbf{y}, \end{aligned} \tag{6}$$

and

$$S(\tau_\varepsilon; \mathbf{y}) = \frac{\partial \ell^R(\boldsymbol{\tau}; \mathbf{y})}{\partial \tau_\varepsilon}$$

$$
\begin{aligned}
&= -\frac{1}{2}\text{tr}(\mathbf{V}^{-1}\mathbf{I}_n) + \frac{1}{2}\text{tr}\big[(\mathbf{X}^T\mathbf{V}^{-1}\mathbf{X})^{-1}\mathbf{X}^T\mathbf{V}^{-1}\mathbf{I}_n\mathbf{V}^{-1}\mathbf{X}\big] + \frac{1}{2}\mathbf{y}'\mathbf{P}\mathbf{I}_n\mathbf{P}\mathbf{y} \\
&= -\frac{1}{2}\text{tr}\Big\{\big[\mathbf{V}^{-1} - \mathbf{V}^{-1}\mathbf{X}(\mathbf{X}^T\mathbf{V}^{-1}\mathbf{X})^{-1}\mathbf{X}^T\mathbf{V}^{-1}\big]\mathbf{I}_n\Big\} + \frac{1}{2}\mathbf{y}'\mathbf{P}\mathbf{I}_n\mathbf{P}\mathbf{y} \\
&= -\frac{1}{2}\text{tr}(\mathbf{P}\mathbf{I}_n) + \frac{1}{2}\mathbf{y}'\mathbf{P}\mathbf{I}_n\mathbf{P}\mathbf{y}. \qquad (7)
\end{aligned}
$$

We further compute the second derivatives of the REML log-likelihood

$$
\begin{aligned}
\frac{\partial^2 \ell^R(\boldsymbol{\tau};\mathbf{y})}{\partial\tau_k\partial\tau_l} &= \frac{1}{2}\text{tr}(\mathbf{P}\mathbf{K}_l\mathbf{P}\mathbf{K}_k) - \frac{1}{2}\mathbf{y}'\mathbf{P}\mathbf{K}_l\mathbf{P}\mathbf{K}_k\mathbf{P}\mathbf{y} - \frac{1}{2}\mathbf{y}'\mathbf{P}\mathbf{K}_k\mathbf{P}\mathbf{K}_l\mathbf{P}\mathbf{y} \\
&= \frac{1}{2}\text{tr}(\mathbf{P}\mathbf{K}_l\mathbf{P}\mathbf{K}_k) - \mathbf{y}'\mathbf{P}\mathbf{K}_k\mathbf{P}\mathbf{K}_l\mathbf{P}\mathbf{y} \\
\frac{\partial^2 \ell^R(\boldsymbol{\tau};\mathbf{y})}{\partial\tau_l\partial\tau_\varepsilon} &= \frac{1}{2}\text{tr}(\mathbf{P}\mathbf{I}_n\mathbf{P}\mathbf{K}_l) - \frac{1}{2}\mathbf{y}'\mathbf{P}\mathbf{I}_n\mathbf{P}\mathbf{K}_l\mathbf{P}\mathbf{y} - \frac{1}{2}\mathbf{y}'\mathbf{P}\mathbf{K}_l\mathbf{P}\mathbf{I}_n\mathbf{P}\mathbf{y} \\
&= \frac{1}{2}\text{tr}(\mathbf{P}\mathbf{I}_n\mathbf{P}\mathbf{K}_l) - \mathbf{y}'\mathbf{P}\mathbf{K}_l\mathbf{P}\mathbf{I}_n\mathbf{P}\mathbf{y} \\
\frac{\partial^2 \ell^R(\boldsymbol{\tau};\mathbf{y})}{\partial\tau_\varepsilon\partial\tau_l} &= \frac{1}{2}\text{tr}(\mathbf{P}\mathbf{K}_l\mathbf{P}\mathbf{I}_n) - \frac{1}{2}\mathbf{y}'\mathbf{P}\mathbf{K}_l\mathbf{P}\mathbf{I}_n\mathbf{P}\mathbf{y} - \frac{1}{2}\mathbf{y}'\mathbf{P}\mathbf{I}_n\mathbf{P}\mathbf{K}_l\mathbf{P}\mathbf{y} \\
&= \frac{1}{2}\text{tr}(\mathbf{P}\mathbf{K}_l\mathbf{P}\mathbf{I}_n) - \mathbf{y}'\mathbf{P}\mathbf{K}_l\mathbf{P}\mathbf{I}_n\mathbf{P}\mathbf{y} \\
\frac{\partial^2 \ell^R(\boldsymbol{\tau};\mathbf{y})}{\partial\tau_\varepsilon^2} &= \frac{1}{2}\text{tr}(\mathbf{P}\mathbf{I}_n\mathbf{P}\mathbf{I}_n) - \frac{1}{2}\mathbf{y}'\mathbf{P}\mathbf{I}_n\mathbf{P}\mathbf{I}_n\mathbf{P}\mathbf{y} - \frac{1}{2}\mathbf{y}'\mathbf{P}\mathbf{I}_n\mathbf{P}\mathbf{I}_n\mathbf{P}\mathbf{y} \\
&= \frac{1}{2}\text{tr}(\mathbf{P}\mathbf{I}_n\mathbf{P}\mathbf{I}_n) - \mathbf{y}'\mathbf{P}\mathbf{I}_n\mathbf{P}\mathbf{I}_n\mathbf{P}\mathbf{y},
\end{aligned}
$$

where $k, l \in \mathcal{L}$.

Since we have

$$
\begin{aligned}
\mathbb{E}(\mathbf{y}'\mathbf{P}\mathbf{K}_k\mathbf{P}\mathbf{K}_l\mathbf{P}\mathbf{y}) &= \text{tr}(\mathbf{P}\mathbf{K}_l\mathbf{P}\mathbf{K}_k) \\
\mathbb{E}(\mathbf{y}'\mathbf{P}\mathbf{K}_k\mathbf{P}\mathbf{I}_n\mathbf{P}\mathbf{y}) &= \text{tr}(\mathbf{P}\mathbf{I}_n\mathbf{P}\mathbf{K}_k) \\
\mathbb{E}(\mathbf{y}'\mathbf{P}\mathbf{I}_n\mathbf{P}\mathbf{I}_n\mathbf{P}\mathbf{y}) &= \text{tr}(\mathbf{P}\mathbf{I}_n\mathbf{P}\mathbf{I}_n),
\end{aligned}
$$

then the average information matrix is given by

$$
\mathbf{AI}(\boldsymbol{\theta};\mathbf{y}) \qquad (8)
$$

$$
= \begin{bmatrix}
\frac{1}{2}\Big[\frac{\partial^2\ell^R(\boldsymbol{\tau};\mathbf{y})}{\partial\tau_s^2} + \mathbb{E}\big(\frac{\partial^2\ell^R(\boldsymbol{\tau};\mathbf{y})}{\partial\tau_s^2}\big)\Big] & \frac{1}{2}\Big[\frac{\partial^2\ell^R(\boldsymbol{\tau};\mathbf{y})}{\partial\tau_s\partial\tau_t} + \mathbb{E}\big(\frac{\partial^2\ell^R(\boldsymbol{\tau};\mathbf{y})}{\partial\tau_s\partial\tau_t}\big)\Big] & \frac{1}{2}\Big[\frac{\partial^2\ell^R(\boldsymbol{\tau};\mathbf{y})}{\partial\tau_s\partial\tau_\varepsilon} + \mathbb{E}\big(\frac{\partial^2\ell^R(\boldsymbol{\tau};\mathbf{y})}{\partial\tau_s\partial\tau_\varepsilon}\big)\Big] \\
\frac{1}{2}\Big[\frac{\partial^2\ell^R(\boldsymbol{\tau};\mathbf{y})}{\partial\tau_t\partial\tau_s} + \mathbb{E}\big(\frac{\partial^2\ell^R(\boldsymbol{\tau};\mathbf{y})}{\partial\tau_t\partial\tau_s}\big)\Big] & \frac{1}{2}\Big[\frac{\partial^2\ell^R(\boldsymbol{\tau};\mathbf{y})}{\partial\tau_t^2} + \mathbb{E}\big(\frac{\partial^2\ell^R(\boldsymbol{\tau};\mathbf{y})}{\partial\tau_t^2}\big)\Big] & \frac{1}{2}\Big[\frac{\partial^2\ell^R(\boldsymbol{\tau};\mathbf{y})}{\partial\tau_t\partial\tau_\varepsilon} + \mathbb{E}\big(\frac{\partial^2\ell^R(\boldsymbol{\tau};\mathbf{y})}{\partial\tau_t\partial\tau_\varepsilon}\big)\Big] \\
\frac{1}{2}\Big[\frac{\partial^2\ell^R(\boldsymbol{\tau};\mathbf{y})}{\partial\tau_\varepsilon\partial\tau_s} + \mathbb{E}\big(\frac{\partial^2\ell^R(\boldsymbol{\tau};\mathbf{y})}{\partial\tau_\varepsilon\partial\tau_s}\big)\Big] & \frac{1}{2}\Big[\frac{\partial^2\ell^R(\boldsymbol{\tau};\mathbf{y})}{\partial\tau_\varepsilon\partial\tau_t} + \mathbb{E}\big(\frac{\partial^2\ell^R(\boldsymbol{\tau};\mathbf{y})}{\partial\tau_\varepsilon\partial\tau_t}\big)\Big] & \frac{1}{2}\Big[\frac{\partial^2\ell^R(\boldsymbol{\tau};\mathbf{y})}{\partial\tau_\varepsilon^2} + \mathbb{E}\big(\frac{\partial^2\ell^R(\boldsymbol{\sigma};\mathbf{y})}{\partial\tau_\varepsilon^2}\big)\Big]
\end{bmatrix}
$$

$$
= \frac{1}{2}\begin{bmatrix}
\mathbf{y}'\mathbf{P}\mathbf{K}_s\mathbf{P}\mathbf{K}_s\mathbf{P}\mathbf{y} & \mathbf{y}'\mathbf{P}\mathbf{K}_s\mathbf{P}\mathbf{K}_t\mathbf{P}\mathbf{y} & \mathbf{y}'\mathbf{P}\mathbf{K}_s\mathbf{P}\mathbf{I}_n\mathbf{P}\mathbf{y} \\
\mathbf{y}'\mathbf{P}\mathbf{K}_t\mathbf{P}\mathbf{K}_s\mathbf{P}\mathbf{y} & \mathbf{y}'\mathbf{P}\mathbf{K}_t\mathbf{P}\mathbf{K}_t\mathbf{P}\mathbf{y} & \mathbf{y}'\mathbf{P}\mathbf{K}_t\mathbf{P}\mathbf{I}_n\mathbf{P}\mathbf{y} \\
\mathbf{y}'\mathbf{P}\mathbf{I}_n\mathbf{P}\mathbf{K}_s\mathbf{P}\mathbf{y} & \mathbf{y}'\mathbf{P}\mathbf{I}_n\mathbf{P}\mathbf{K}_t\mathbf{P}\mathbf{y} & \mathbf{y}'\mathbf{P}\mathbf{I}_n\mathbf{P}\mathbf{I}_n\mathbf{P}\mathbf{y}
\end{bmatrix}. \qquad (9)
$$

We can apply the Newton's method for the estimates of variance components $\boldsymbol{\phi}$ by the following iterative update

$$\boldsymbol{\tau}^{(t+1)} = \boldsymbol{\tau}^{(t)} + \mathbf{AI}(\boldsymbol{\theta}^{(t)}; \mathbf{y})\mathbf{S}(\boldsymbol{\tau}^{(t)}; \mathbf{y}), \tag{10}$$

and further obtain the estimates of $\boldsymbol{\beta}$. In practice, we implement the "lmm_aireml" function in the R package 'gaston' to refine the estimates more efficiently.

#### 1.2 Test 1: Overall Test To Identify uTSVG

To detect TSVGs, we consider the following null and alternative hypotheses

$$\begin{cases} H_0^{(1)} : \tau_t = \tau_s = 0 \\ H_1^{(1)} : \tau_l \neq 0, \quad l \in \mathcal{L} = \{s, t\} \end{cases} \tag{11}$$

Under the null hypothesis $H_0^{(1)}$, the null model is given by

$$\mathbf{y} \sim MVN\left(\mathbf{X}\beta, \mathbf{V_0}\right), \tag{12}$$

where $\mathbf{V_0} = \tau_\varepsilon \mathbf{I}_n$.

Then, by fitting the above null model, we obtain $\mathbf{P} = \frac{1}{\tau_\varepsilon}(\mathbf{I}_n - \mathbf{X}(\mathbf{X}'\mathbf{X})^{-1}\mathbf{X}')$ and the estimation of $\hat{\beta} = (\mathbf{X}'\mathbf{X})^{-1}\mathbf{X}'\mathbf{y}$ and $\hat{\tau}_\varepsilon = \frac{1}{n-\nu}(\mathbf{y} - \mathbf{X}\hat{\beta})'(\mathbf{y} - \mathbf{X}\hat{\beta})$.

We use the score test statistics

$$Q(\mathbf{y}) = \mathbf{y}'\mathbf{P} \textstyle\sum_{l \in \mathcal{L}} \mathbf{K}_l \mathbf{P}\mathbf{y}, \tag{13}$$

which follows a mixture of chi-square distribution under the null,

$$Q(\mathbf{y})/\tau_\varepsilon \sim \sum_{i=1}^{J} \lambda_i \chi_1^2$$

where $\{\lambda_i\}$ are eigenvalues of $\mathbf{P} \sum_{l \in \mathcal{L}} \mathbf{K}_l \mathbf{P}$, $\chi_1^2$ is a random variable independently and identically distributed from a chi-square distribution with the degree of freedom being one, and $J$ is the number of non-zero eigen values. In practice, $\tau_\varepsilon$ is unknown, so we replace it with its restricted maximum likelihood(REML) estimate $\hat{\tau}_\epsilon$. We apply the score test [1] to calculate the distribution of $Q(\mathbf{y})/\hat{\tau}_\varepsilon$ using the Davies method.

#### 1.3 Test 2: Individual Test To Assess The Spatial or Temporal Effect

For the testing of conditional individual effect, for each of $\ell \in \mathcal{L} = \{s, t\}$, we consider the null $H_0^{(2)} : \tau_\ell = 0$.

$$\begin{cases} H_0^{(2)} : \tau_l = 0 \\ H_1^{(2)} : \tau_l \neq 0 \end{cases} \tag{14}$$

To simplify the notation, let $-\ell = \{s,t\}\backslash\ell$. Therefore, under null $H_0^{(2)}$ model, $\mathrm{Var}(\mathbf{y}) = \mathbf{V}_{-\ell} + \tau_\varepsilon \mathbf{K}_\varepsilon = \tau_{-\ell}K_{-\ell} + \tau_\varepsilon\mathbf{K}_\varepsilon$

The score function is

$$\tfrac{\partial \ell_R(\boldsymbol{\theta})}{\partial \tau_\ell} = -\tfrac{1}{2}\,\mathrm{tr}\left(\boldsymbol{K}_\ell \boldsymbol{P}_{-\ell}\right) + \tfrac{1}{2}\boldsymbol{y}'\boldsymbol{P}_{-\ell}\boldsymbol{K}_\ell\boldsymbol{P}_{-\ell}\boldsymbol{y}$$

where $\boldsymbol{P}_{-\ell} = \boldsymbol{V}_\ell^{-1} - \boldsymbol{V}_{-\ell}^{-1}\boldsymbol{X}\left(\boldsymbol{X}'\boldsymbol{V}_{-\ell}^{-1}\boldsymbol{X}\right)^{-1}\boldsymbol{X}'\boldsymbol{V}_{-\ell}^{-1}$ is the projection matrix under the null.

Then, the test statistic is given by $U_1(\mathbf{y}) = \frac{1}{2}\mathbf{y}'\mathbf{P}_{-\ell}\mathbf{K}_\ell\mathbf{P}_{-\ell}\mathbf{y}$ and $U_1(\mathbf{y}) \sim \kappa\chi^2_\nu$, under the null hypothesis.

We use the Satterthwaite method to approximate the distribution of $\kappa\chi^2_\nu$, where the scale parameter $\kappa$ and degree of freedom $\nu$ are estimated by Method of Moments (MOM) by equating the mean and variance of the test statistics with the distribution. Because $U_1(\mathbf{y})$ is a quadratic term of $\mathbf{y}$, we can get its mean and variance easily.

$$\begin{cases} \delta = \kappa\nu = \mathbb{E}(U_1(\mathbf{y})) \\ \quad = \dfrac{1}{2}\left[\mathrm{tr}(\mathbf{P}_{-\ell}\mathbf{K}_\ell\mathbf{P}_{-\ell}\mathbf{V}_{-\ell}) + \beta'\mathbf{X}'\mathbf{P}_{-\ell}\mathbf{K}_\ell\mathbf{P}_{-\ell}\mathbf{X}\beta\right] \\ \quad = \dfrac{1}{2}\,\mathrm{tr}(\mathbf{K}_\ell\mathbf{P}_{-\ell}\mathbf{V}_{-\ell}\mathbf{P}_{-\ell}) = \dfrac{1}{2}\,\mathrm{tr}(\mathbf{K}_\ell\mathbf{P}_{-\ell}), \\ \gamma = 2\kappa^2\nu = \mathrm{Var}(U_1(\mathbf{y})) \\ \quad = \dfrac{1}{2}\,\mathrm{tr}(\mathbf{P}_{-\ell}\mathbf{K}_\ell\mathbf{P}_{-\ell}\mathbf{V}_{-\ell}\mathbf{P}_{-\ell}\mathbf{K}_\ell\mathbf{P}_{-\ell}\mathbf{V}_{-\ell}) \\ \qquad + \beta'\mathbf{X}'\mathbf{P}_{-\ell}\mathbf{K}_\ell\mathbf{P}_{-\ell}\mathbf{V}_{-\ell}\mathbf{P}_{-\ell}\mathbf{K}_\ell\mathbf{P}_{-\ell}\mathbf{X}\beta \\ \quad = \dfrac{1}{2}\,\mathrm{tr}(\mathbf{K}_\ell\mathbf{P}_{-\ell}\mathbf{K}_\ell\mathbf{P}_{-\ell}) \end{cases} \tag{15}$$

In practice, we do not know the true value of $\tau_{-\ell}$ and usually replace them by the its MLE under the null model. Therefore, the notation after plug-in is $\hat{\mathbf{P}}_{-\ell} = \hat{\boldsymbol{V}}_\ell^{-1} - \hat{\boldsymbol{V}}_{-\ell}^{-1}\boldsymbol{X}\left(\boldsymbol{X}'\hat{\boldsymbol{V}}_{-\ell}^{-1}\boldsymbol{X}\right)^{-1}\boldsymbol{X}'\hat{\boldsymbol{V}}_{-\ell}^{-1}$, where $\hat{\mathbf{V}}_{-\ell} = \tau_{-\ell}K_{-\ell}$

To account for this substitution, we replace $\tilde{\mathcal{I}}$ by $\tilde{\mathcal{I}}_{\ell,\ell} = \mathcal{I}_{\ell,\ell} - \mathcal{I}_{\ell,-\ell}\mathcal{I}_{-\ell,-\ell}{}^{-1}\mathcal{I}_{-\ell,\ell}$. The Fisher information matrix is given by

$$\mathcal{I}(\boldsymbol{\tau}_{-\ell}, \tau_\ell) = \begin{bmatrix} \mathcal{I}_{-\ell,-\ell} & \mathcal{I}_{-\ell,\ell} \\ \mathcal{I}_{\ell,-\ell} & \mathcal{I}_{\ell,\ell} \end{bmatrix},$$

$$\mathcal{I}_{-\ell,-\ell} = \frac{1}{2}\begin{bmatrix} \mathrm{tr}(\mathbf{P}_{-\ell}\mathbf{P}_{-\ell}) & \mathrm{tr}(\mathbf{P}_{-\ell}\mathbf{P}_{-\ell}\mathbf{K}_1) & \ldots & \mathrm{tr}(\mathbf{P}_{-\ell}\mathbf{P}_{-\ell}\mathbf{K}_L) \\ \mathrm{tr}(\mathbf{P}_{-\ell}\mathbf{K}_1\mathbf{P}_{-\ell}) & \mathrm{tr}(\mathbf{P}_{-\ell}\mathbf{K}_1\mathbf{P}_{-\ell}\mathbf{K}_2) & \ldots & \mathrm{tr}(\mathbf{P}_{-\ell}\mathbf{K}_1\mathbf{P}_{-\ell}\mathbf{K}_L) \\ \vdots & \vdots & \vdots & \vdots \\ \mathrm{tr}(\mathbf{P}_{-\ell}\mathbf{K}_L\mathbf{P}_{-\ell}) & \mathrm{tr}(\mathbf{P}_{-\ell}\mathbf{K}_L\mathbf{P}_{-\ell}\mathbf{K}_2) & \ldots & \mathrm{tr}(\mathbf{P}_{-\ell}\mathbf{K}_L\mathbf{P}_{-\ell}\mathbf{K}_L) \end{bmatrix},$$

$$\mathcal{I}_{-\ell,\ell} = \frac{1}{2}\left[\mathrm{tr}\left(\mathbf{P}_{-\ell}\mathbf{P}_{-\ell}\mathbf{K}_\ell\right), \mathrm{tr}\left(\mathbf{P}_{-\ell}\mathbf{K}_1\mathbf{P}_{-\ell}\mathbf{K}_\ell\right), \ldots, \mathrm{tr}\left(\mathbf{P}_{-\ell}\mathbf{K}_m\mathbf{P}_{-\ell}\mathbf{K}_\ell\right)\right]',$$

$$\mathcal{I}_{\ell,\ell} = \frac{1}{2}\,\mathrm{tr}\left(\mathbf{P}_{-\ell}\mathbf{K}_\ell\mathbf{P}_{-\ell}\mathbf{K}_\ell\right).$$

Therefore, $\tilde{\gamma} = \tilde{\mathcal{I}}_{\ell,\ell}$ , $\tilde{\delta} = \frac{1}{2}\,\mathrm{tr}(\mathbf{K}_\ell \hat{\mathbf{P}}_{-\ell})$, $\hat{\kappa} = \frac{\tilde{\gamma}}{2\tilde{\delta}}$ and $\hat{\nu} = \frac{2\tilde{\delta}^2}{\tilde{\gamma}}$

##### 1.4 Kernel Matrix

The kernel matrix, which represents the spatial correlation pattern and temporal correlation pattern of spots within the target tissue, is crucial for parameter estimation and statistical inference. In this work, we employ Gaussian kernel for both spatial and temporal correlation, defined as $\boldsymbol{K}_s(\boldsymbol{s_i}, \boldsymbol{s_j}) = \exp(-\frac{\|\boldsymbol{s_i}-\boldsymbol{s_j}\|^2}{h})$, where $\|\boldsymbol{s_i} - \boldsymbol{s_j}\|^2$ denotes the Euclidean distance between spots $\boldsymbol{s_i} = (s_{i,1}, s_{i,2})$ and $\boldsymbol{s_2} = (s_{j,1}, s_{j,2})$, and $h$ denotes the bandwidth. The Euclidean distance is spatial rotation invariant, therefore, TESSA also inherits this good property. That is to say, even if the tissue is rotated when ST platform measures its gene expression, the result of TESSA is consistent with spatial coordinates rotations.

To determine the appropriate bandwidth $h$, we adhere to the method proposed by spatialPCA[2]. Specifically, for datasets with 5000 spots or fewer, non-parametric Sheather-Jones' bandwidths[3] are computed for each gene, and the median of these values is used as the common bandwidth $h$; for datasets with more than 5000 spots, Silverman's rule of thumb [4] are computed for each gene, and the median of these values is used as the common bandwidth $h$. The user can also apply the cosine kernel with different parameters and get different p-values, then these p-values can be combined with the Cauchy combination rule as implemented in SPARK[5].

#### 2 Simulation Results In Normal Scenario

The first set of simulations evaluated the performance of the TESSA stage 1 overall test for uTSVG detection. We compared it with SPARK for SVG detection and with DataThin for double-dipping correction. In this section, we focus on the Normal data generation scenario, as it reveals the efficacy of the tests in terms of both false-positive control and statistical power under ideal conditions where the data distribution matches the model assumptions.

**Assess the overall null:** Under the null setting with only noise genes, those without spatial or temporal patterns, TESSA-oracle effectively controlled the type I error. In contrast, with the estimated pseudotime, TESSA-slingshot exhibited inflated false positives due to double dipping. Both TESSA-slingshot-LOO and TESSA-slingshot-datathin successfully controlled type I error (Fig.S1.d, left panel). This validates that DataThin performs well when model assumptions are satisfied. The results also demonstrate that LOO provides a model-free correction for double dipping, since its mechanism of removing the tested gene from pseudotime estimation is independent of the data generation process. As SPARK relies solely on spatial location information, it is unaffected by the double-dipping issue.

**Assess the overall power:** Under the alternative setting with a mixture of uTSVGs and noise genes, TESSA-oracle, TESSA-slingshot, and TESSA-slingshot-LOO achieved comparable accuracy in distinguishing uTSVGs from noise genes. In the ideal Normal scenario, the higher power of TESSA-slingshot offsets the inflation of false positives from double dipping, resulting in overall accuracy comparable to TESSA-slingshot-LOO. By contrast, the power loss of DataThin is consistent across both the Normal and NB scenarios and remains worse than that of TESSA-slingshot-LOO.

**Assess the individual null:** The second set of simulations evaluated the performance of the TESSA stage 2 individual test for detecting SVGs and TVGs separately. For TVG detection, we compared against DataThin as a double-dipping correction method, while for SVG detection there is no double-dipping issue.

Under the null setting with only noise genes, TESSA-oracle effectively controlled the type I error when testing TVGs. Consistent with the first set of simulations, TESSA-slingshot exhibited inflated false positives due to double dipping. Both TESSA-slingshot-LOO and TESSA-slingshot-datathin successfully controlled type I error (Fig.S1.d, left panel).

**Assess the individual power:** Under the alternative setting with a mixture of uTSVGs and noise genes, TESSA-oracle and TESSA-slingshot achieved similar accuracy when testing for SVGs. For TVG detection, TESSA-oracle, which uses the ground-truth pseudotime, attained the highest accuracy in distinguishing uTSVGs from noise genes. TESSA-slingshot and TESSA-slingshot-LOO showed varying degrees of deviation, but when the signal strength was large, all three methods demonstrated comparable accuracy (Fig.S1.e,f). By contrast, the power loss of DataThin remained substantial compared to TESSA-slingshot-LOO.

### 3 Additional Details of Real Data Analysis

#### 3.1 Pseudotime Estimation

We used slingshot[6] to estimate pseudotime for all three datasets. We opted for slingshot rather than spatial pseudotime methods to avoid confounding temporal and spatial signals, since spatial methods incorporate spatial coordinates into pseudotime construction. This consideration is especially important in heterogeneous tissues such as tumors, where spatial methods like SpaceFlow may over-smooth sharp boundaries, potentially obscuring biological variations that are enriched along pseudotime (Fig. S4). In practice, users may extend our framework to other pseudotime estimation methods. Signature gene selection is also critical for pseudotime estimation; we recommend using strong and reliable clinical markers or signature genes derived from scRNA-seq, as they provide meaningful prior knowledge.

The standard slingshot pipeline designed for scRNA-seq datasets includes data normalization, feature selection, principal component analysis (PCA), clustering, specification of a starting cluster, and pseudotime estimation. We modified the feature selection step by using a pre-defined signature gene set $\mathcal{M}$, including known cell type markers and tumor state signature genes.

For the human pancreatic cancer data[7], available through dbGaP under accession number PRJNA1124001 with image files at https://zenodo.org/records/13379726, we used sample S2A as an example. Following the slingshot procedure, we selected 31 signature genes related to pancreatic tumor progression. These include markers for normal ductal cells (*EPCAM*, *GATA6*, *CFTR*, *KRT7*, *CDH1*, *HNF1A*, *FOXA2*), tumor ductal cells (*MUC1*, *MUC5AC*, *S100P*, *SOX9*, *ZEB1*, *SNAI2*, *TP63*, *FN1*, *CD44*,

*VIM*, *TFF1*, *TFF2*), and stromal cancer-associated fibroblasts (*FN1*, *SPARC*, *ACTA2*, *PDGFRB*, *POSTN*, *LOXL2*, *MMP2*, *MMP9*, *FAP*, *TGFB1*, *CXCL12*, *CCL2*). We used Seurat to perform preprocessing steps including SCTransform normalization, PCA, and Louvain clustering. We set cluster 4, corresponding to the PanIN region (validated by PanIN markers in Fig.6c), as the starting cluster. Slingshot produced an estimated pseudotime with two lineages (Fig.S2): lineage 1 captures PDAC tumor progression, while lineage 2 describes trajectories among PanIN, ductal, and acinar cells.

For the human lung cancer dataset (GEO accession GSE189487), we used sample TD8. We extracted the top 30 level-3 cell type signature genes from the scRNA-seq Human Lung Cell Atlas (HLCA)[8], which is available from cellxgene (https://cellxgene.cziscience.com/collections/6f6d381a-7701-4781-935c-db10d30de293 ). Using Seurat::FindAllMarkers(), we identified level-3 cell type markers and filtered by pct.1 $> 0.3$, then selected the top 50 genes with the largest average $\log_2$ fold change among the 200 most significant genes. These representative markers were used as signature genes for pseudotime inference. Following the standard slingshot pipeline, we used SCTransform normalization, PCA, and Louvain clustering. Cluster 0, corresponding to the lung adenocarcinoma region (validated by MUC1 expression in Fig.7c), was set as the starting cluster. Slingshot inferred three lineages (Fig.S6): lineage 1 reflects developmental transitions between LUAD and normal epithelium, lineage 2 captures transitions between LUAD and stroma, and lineage 3 captures LUAD–lymph interactions.

For the mouse primary visual cortex data[9], available at https://github.com/zhengli09/BASS-Analysis/tree/master/data , we analyzed samples BZ5, BZ9, and BZ14. Inspired by 15 well-known marker genes, we constructed pseudotime trajectories across excitatory neurons (*Slc17a7*, *Cux2*, *Bcl6*, *Nptx2*, *Rorb*, *Tbr1*), inhibitory neurons (*Pvalb*, *Sst*, *Vip*), astrocytes (*Aqp4*), oligodendrocytes (*Mog*, *Pdgfra*), microglia (*Itgam*), and developmental markers (*Foxp2*, *Pax6*). Seurat preprocessing included normalization with regression on sample ID using SCTransform, and PCA embedding based on these signature genes. With the cluster number set to one, slingshot automatically identified the starting point, resulting in a single lineage pseudotime trajectory.

For the human dorsolateral prefrontal cortex data[10], available at https://github.com/LieberInstitute/HumanPilot , we analyzed samples 151507, 151508, 151509, and 151510. We selected 28 signature genes representing major cortical cell types: excitatory neurons (*SLC17A7*, *CUX2*, *BCL6*, *NPTX2*, *RORB*, *TBR1*), inhibitory neurons (*PVALB*, *SST*, *VIP*, *NPY*, *NOS1*, *CRH*, *CCK*, *RELN*, *LHX6*), astrocytes (*AQP4*), oligodendrocytes (*MOG*, *PDGFRA*), microglia (*ITGAM*), neuronal activity markers (*FOS*, *FOSB*, *EGR1*, *EGR2*, *EGR4*, *NPAS4*, *ARC*), and developmental markers (*FOXP2*, *PAX6*). Preprocessing with Seurat included normalization using SCTransform (regressing out sample ID) and PCA embedding restricted to these signature genes. With the cluster number set to one, slingshot automatically selected the starting point, yielding a single lineage pseudotime trajectory.

#### 3.2 Domain Detection Analysis

We initially applied two domain detection algorithms to all datasets. For the 10x Visium cancer datasets, we selected SeuratPCA as the primary method to highlight differences between our uTSVGs and SPARK SVGs. For the Starmap mouse cortex dataset, we employed the multi-sample extension of SpatialPCA as the main algorithm. Results from the alternative domain detection approach are provided in the supplementary figures.

SeuratPCA takes the gene expression matrix as input, performs principal component analysis (PCA) to reduce dimensionality, and applies the Louvain clustering algorithm to identify spatial domains. To improve accuracy, we further refined the domain labels using SpatialPCA::refine_cluster_10x().

SpatialPCA uses both gene expression and spatial location as input, applies probabilistic PCA to reduce the data into low-dimensional spatial PCs, and then employs the walktrap clustering algorithm to detect spatial domains. The final clustering results were also refined using SpatialPCA::refine_cluster_10x(). In the multi-sample extension of SpatialPCA, the default setting uses common SPARK SVGs across samples as input features, constructs a block-diagonal location kernel from spatial coordinates of each sample, and then follows the same downstream steps.

#### 3.3 Signature Scores Calculation

The signature scores is calculated as $\log\left(\frac{1}{|\mathcal{G}|}\sum_{g\in\mathcal{G}}\left(e^{X_{g,\cdot}}-1\right)+1\right)$ for the signature gene set $\mathcal{G}$. TLS signature score is based on gene set *CETP*, *CCR7*, *SELL*, *LAMP3*, *CCL19*, *CXCL10*, *CXCL11*, *CXCL13*, *CCL21*. CAF signature score is based on gene set *LUM*, *COL1A1*, *COL3A1*, *COL4A1*, *MMP2*, *FAP*, *S100A4*, *PDGFRA*, *PDPN*, *ACTA2*, *MFGE8*. Smooth Muscle signature score is based on gene set *ACTA2*, *CCN5*, *DES*, *MYH11*, *TAGLN*.

#### 3.4 Gene Set Enrichment Analysis

The gene set enrichment analyses (GSEA) were conducted using the `gProfiler2` package. We employed the `gost` function to assess the significance of pathways associated with TESSA TVGs and SVGs, based on GO, KEGG, and Reactome gene sets. Specifically, we used the default "g_SCS" algorithm for multiple testing correction and set the significance threshold at 0.05.

For the human pancreatic cancer dataset, we first performed GSEA using lineage-specific TVGs and SVGs. For lineage 1, we analyzed 1,486 unique TVGs and 1,468 unique SVGs; for lineage 2, we analyzed 1,013 unique TVGs and 1,963 unique SVGs. Here, unique TVGs refer to genes identified as TVGs but not SVGs, while unique SVGs refer to genes identified as SVGs but not TVGs.

Next, we performed GSEA using TVGs and SVGs that were also differentially expressed (DE) in domain 10. DE genes were identified with `Seurat::FindAllMarkers()`, filtered using pct.1 $>$ 0.6 and pct.2 $<$ 0.3 to focus on genes specifically expressed in domain 10, such as those shown in Fig.6e. We then intersected TVGs with domain 10 DE genes as the input for GSEA (Fig.6f, left panel) and intersected SVGs with domain 10 DE genes as the input for GSEA (Fig. 6f, right panel).

For the human lung adenocarcinoma dataset, we similarly performed GSEA using lineage-specific TVGs and SVGs. Given the substantial overlap of genes across the three lineages, we restricted the analysis to TVGs and SVGs uniquely significant to one lineage. For lineage 1, we analyzed 2,135 unique TVGs and 165 SVGs; for lineage 2, 78 unique TVGs and 182 SVGs; and for lineage 3, 114 unique TVGs and 180 SVGs.

#### 3.5 Computational Time

For the PDAC S2A dataset, which contains 11,115 genes, 3,142 spots, and 2 lineages, we evaluated the computational performance of various methods on a desktop with an Apple M1 Pro CPU and 16 GB of RAM. The complete TESSA stage 1 overall test finished in approximately 19.2 minutes. The stage 1 test without the leave-one-out (LOO) component required about 212.53 seconds for all genes across both lineages. On average, the LOO correction for each double-dipping gene took approximately 15.7 seconds per lineage. The TESSA stage 2 individual test required about 46 seconds per gene per lineage, whether for the TVG or SVG test.

### 4 Supplementary Figures

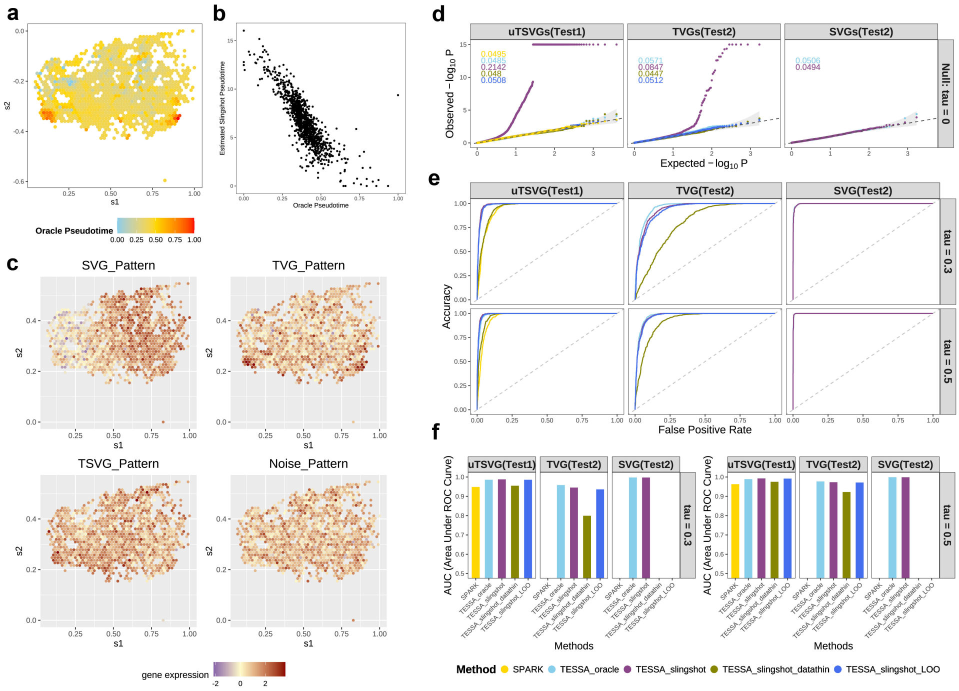

**Fig. S1**: **Simulation data generation and results under the Normal scenario.** **a**. Spatial distribution of the oracle pseudotime. **b**. Scatter plot comparing oracle pseudotime with pseudotime estimated by Slingshot. **c**. Spatial visualization of normalized gene expression patterns for representative genes corresponding to three signal patterns and a noise pattern. **d**. Results of both stage 1 and stage 2 tests under the null case. **e**. AUC-ROC curves for uTSVG, TVG, and SVG detection in stage 1 and stage 2 tests under varying signal strengths. **f**. Bar plots of the area under the ROC curve (AUC) for different methods across varying signal strengths, corresponding to the ROC curves shown in (e).

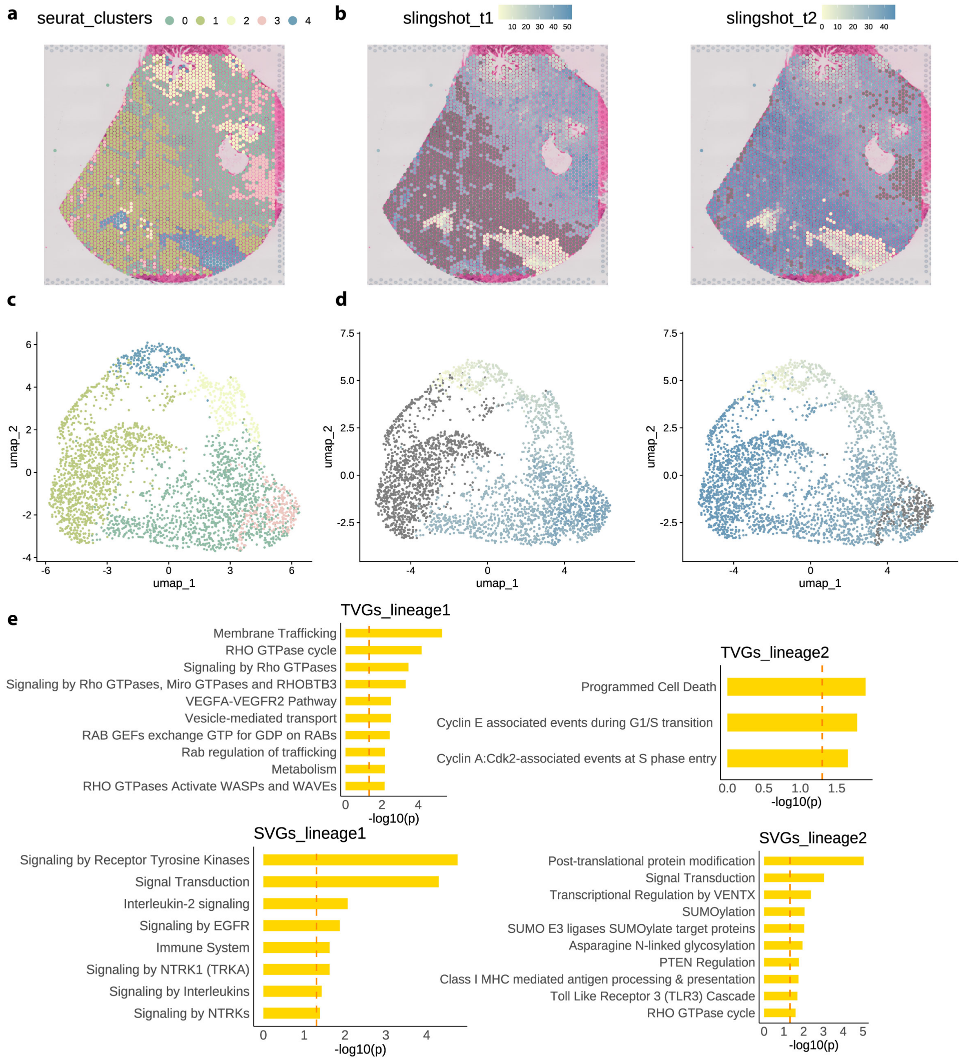

**Fig. S2**: **Slingshot Pseudotime of S2A Pancreatic Cancer Data**. **a**. Visualization of Seurat clusters based on signature genes, mapped onto spatial coordinates. **b**. Visualization of estimated Slingshot pseudotime with spatial coordinates. The slingshot_t1 lineage captures PDAC tumor progression, while slingshot_t2 represents progression among Acinar and Ductal cells. **c**.Visualization of Seurat clusters based on signature genes, projected onto UMAP coordinates. **d**.Visualization of estimated Slingshot pseudotime, projected onto UMAP coordinates. **e**. The Reactome gene set enrichment analysis results pathways of lineage-specific TVGs detected by TESSA stage 2 tests.

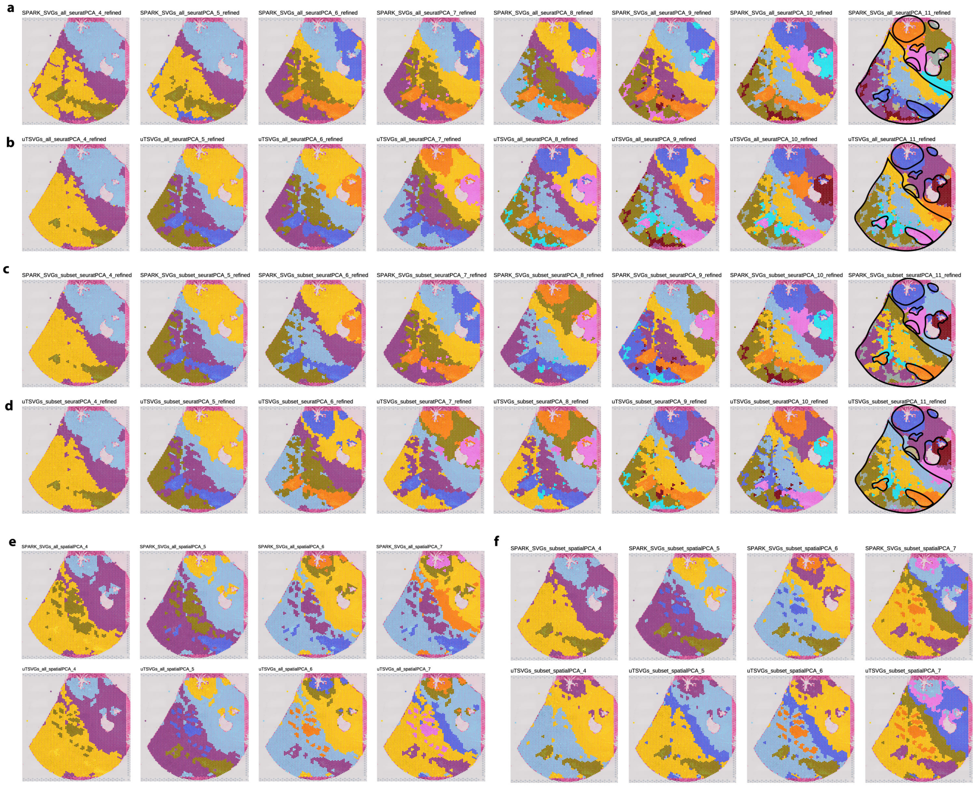

**Fig. S3**: **Domain detection of S2A pancreatic cancer data with varying cluster numbers. a,b**. Spatial domains detected with Seurat using all uTSVGs (**a**) or all SPARK SVGs (**b**) as input, with cluster numbers ranging from 4 to 7. Results were refined with `SpatialPCA::refine_cluster_10x()`. **c,d**. Spatial domains detected with Seurat using the top 2,000 uTSVGs (**c**) or SPARK SVGs (**d**) as input, also refined with `SpatialPCA::refine_cluster_10x()`. **e**. Spatial domains detected with SpatialPCA using all uTSVGs or SPARK SVGs as input (4–7 clusters). **f**. Spatial domains detected with SpatialPCA using the top 2,422 uTSVGs or SPARK SVGs as input (4–7 clusters).

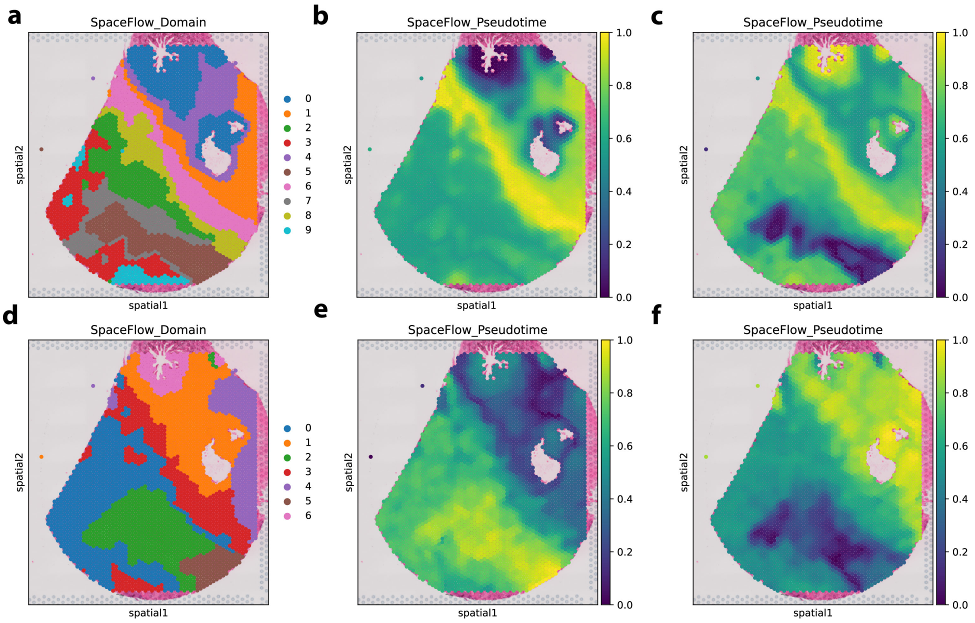

**Fig. S4**: **SpaceFlow pseudotime of S2A pancreatic cancer data.** **a**. Spatial patterns of SpaceFlow clusters derived from 3,000 highly variable genes (HVGs). **b**. Spatial patterns of SpaceFlow pseudotime based on 3,000 HVGs, with the starting point automatically selected by SpaceFlow. The inferred trajectory begins in normal ductal regions and progresses through tumor ductal, acinar, PanIN, tumor-associated fibroblast regions, and finally fibroblast regions adjacent to acinar regions. Notably, some tumor-associated fibroblast regions are temporally close to acinar regions, which is biologically difficult to interpret. **c**. Spatial patterns of SpaceFlow pseudotime based on 3,000 HVGs, with the starting point manually set to a spot in the PanIN region. This choice confuses ductal and fibroblast regions and over-smooths the tumor regions. **d**. Spatial patterns of SpaceFlow clusters derived from the same set of signature genes used for Slingshot pseudotime estimation. With a limited number of signature genes, pseudotime trajectories appear dominated by over-smoothed spatial patterns. **e**. Spatial patterns of SpaceFlow pseudotime based on signature genes, with the starting point automatically selected by SpaceFlow. **f**. Spatial patterns of SpaceFlow pseudotime based on signature genes, with the starting point manually set to a spot in the PanIN region.

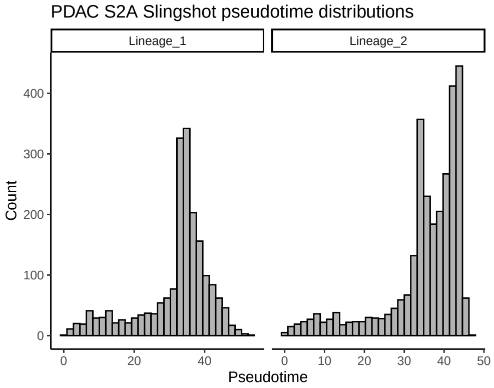

**Fig. S5**: **Histogram of Slingshot Pseudotime on S2A Pancreatic Cancer Data**. The histogram of slingshot pseudotime on S2A sample illustrates that pseudotime might not follow uniform distribution, which motivates us to simulate oracle pseudotime as a mixture of truncated normal distribution in Negative Binomial simulation scenario.

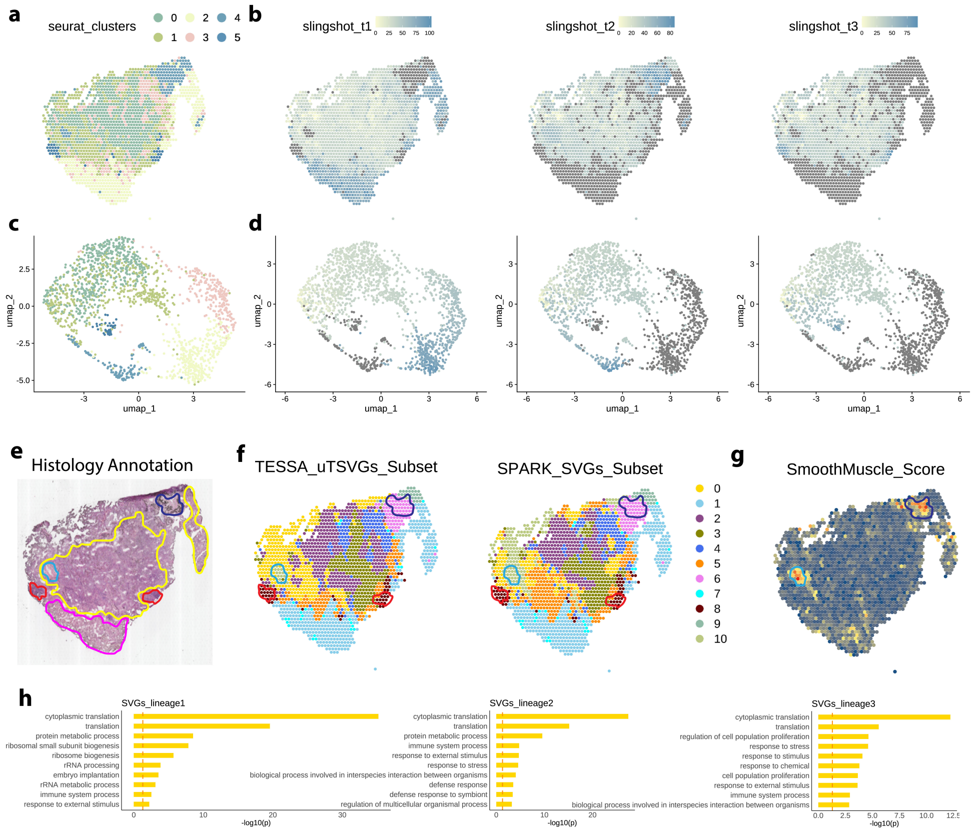

**Fig. S6**: **Slingshot pseudotime of TD8 lung adenocarcinoma data. a**. Seurat clusters based on signature genes mapped onto spatial coordinates. **b**. Estimated Slingshot pseudotime overlaid on spatial coordinates. The `slingshot_t1` lineage captures progression between adenocarcinoma in situ and normal epithelial cells; `slingshot_t2` represents progression between stromal and adenocarcinoma cells; and `slingshot_t3` represents progression between lymphoid and adenocarcinoma cells. **c**. Seurat clusters based on signature genes projected onto UMAP coordinates. **d**. Estimated Slingshot pseudotime projected onto UMAP coordinates. **e**. Histology-based domain annotations as ground truth: yellow, cancer region; red, lymph region; pink, epithelial region; light blue, stromal/cancer vasculature region; dark blue, stromal/normal vasculature region. **f**. Domain detection results showing true and estimated annotations by SeuratPCA. Left: based on TESSA top 1,000 lineage-specific uTSVGs. Right: based on the same number of SPARK SVGs. **g**. Spatial expression patterns of log-normalized smooth muscle signature scores. **h**. GO BP pathway enrichment of three lineage-specific unique SVGs.

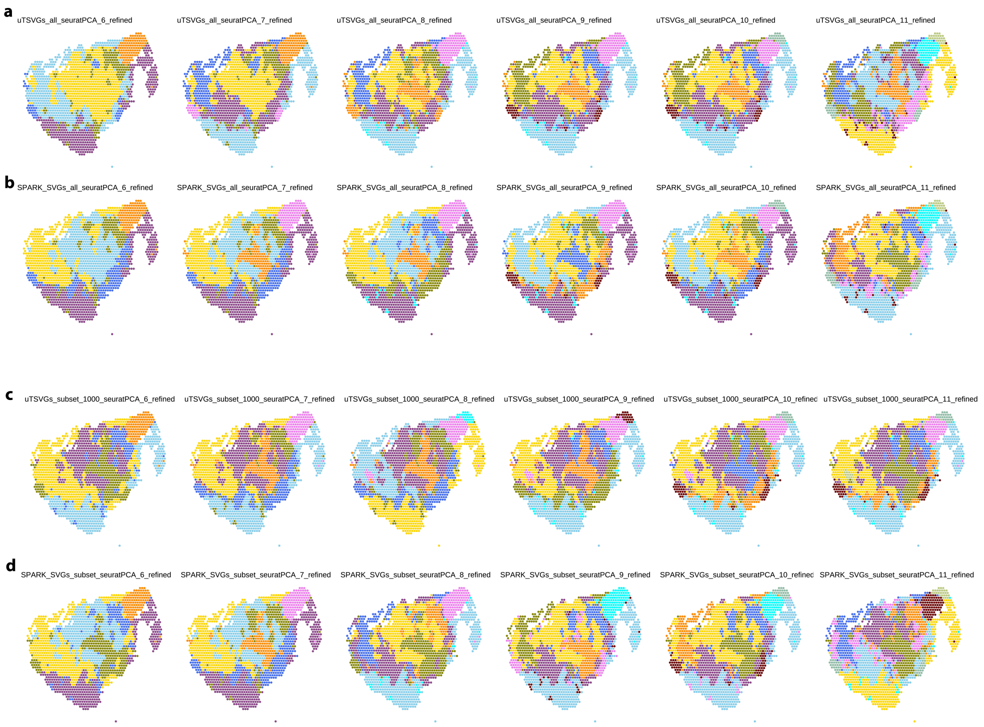

**Fig. S7**: **Seurat Domain Detection of Lung Adenoma Data**. **a,b**. Spatial domain visualization using all uTSVGs(**a**) or all SPARK SVGs(**b**) as input. Domains are detected with refined Seurat, with the number of domains varying from 6 to 12. **c,d**. Spatial domain visualization using top 1,000 lineage-specific uTSVGs(**c**) and the same number of SPARK SVGs(**d**) as input. Domains are detected with refined Seurat, with the number of domains varying from 6 to 12.

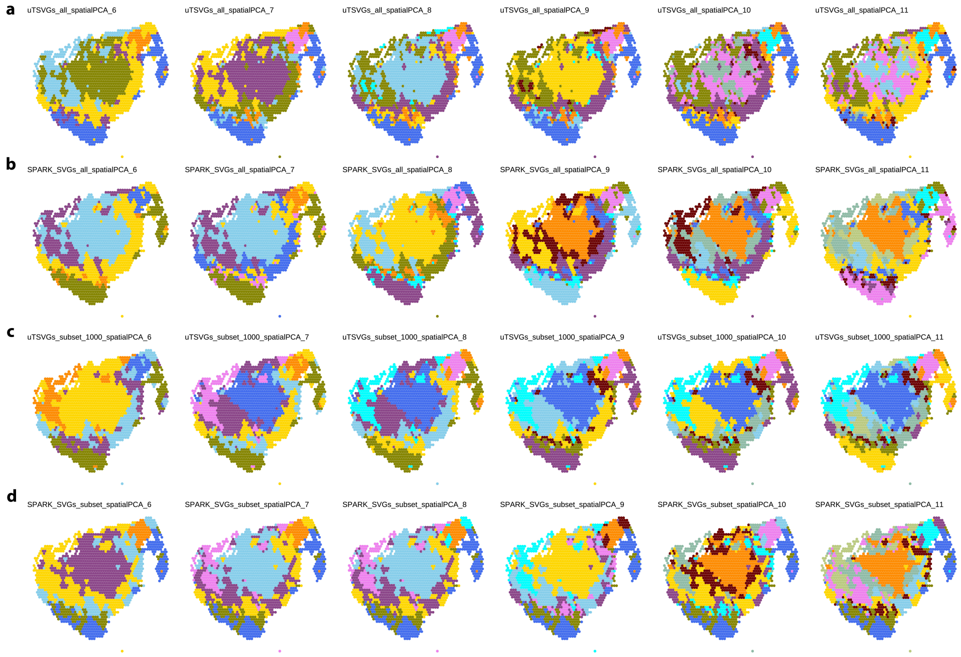

**Fig. S8**: **spatialPCA Domain Detection of Lung Adenoma Data**. **a,b**. Spatial domain visualization using all uTSVGs(**a**) or all SPARK SVGs(**b**) as input. Domains are detected with spatialPCA, with the number of domains varying from 6 to 11. **c,d**. Spatial domain visualization using top 1,000 lineage-specific uTSVGs(**c**) and the same number of SPARK SVGs(**d**) as input. Domains are detected with refined Seurat, with the number of domains varying from 6 to 11.
